# Connectome structure constrains neural activity across scales

**DOI:** 10.64898/2026.11.03.762695

**Authors:** Carter Sale, Michael J. Richardson

**Author notes:** Correspondence: Carter Sale.

## Abstract

Connectomes can be represented at many resolutions, but coarse-graining risks discarding structure relevant to neural dynamics. Here we ask at what resolution a connectome preserves what it predicts about neural activity. In *C. elegans*, direct synaptic connectivity poorly predicts whether stimulating one neuron evokes a response in another. A parameter-free predictor based on paths through the connectome performs substantially better, with most predictive information carried by two-step paths and by the arrangement rather than the number of connections. Coarse-graining neurons into biological cell classes preserves these predictions despite discarding path structure, whereas random partitions degrade them. Connectivity alone identifies alternative coarse-grainings that preserve prediction down to about nine groups corresponding to nervous-system subsystems. Partitions inferred from newly hatched animals preserve adult predictions, and lineage-based partitions in *Drosophila* preserve path structure better than random coarse-graining. Functional prediction can therefore survive at resolutions far coarser than those required to preserve structure.

## 1 Introduction

Connectomes are being reconstructed at ever finer levels of detail, from connections between brain regions down to synapses between individual neurons. For most nervous systems this detail remains out of reach, and connectivity is measured between parcellated regions containing many thousands of cells. The effort required therefore scales steeply with resolution, and it is usually assumed that finer structural detail will yield better accounts of function. Even so, the relationship between neural structure and function has remained difficult to establish.

Testing this relationship is itself difficult because structural and functional measurements are rarely available from the same nervous system at a comparable resolution. *Caenorhabditis elegans*, however, is an exception. Its neurons are individually identifiable, allowing the same cells to be matched across animals and datasets, and its synaptic wiring has been reconstructed at cellular resolution ^1;2;3^. More recently, brain-wide perturbation experiments have stimulated individual neurons while recording responses across the nervous system ^4^. We therefore now have both fine-grained structural information and causal measurements of how activity propa-gates between the same identified cells.

Even with this level of detail, direct synaptic connectivity gives an incomplete account of signal propagation. Many neurons respond to stimulation despite receiving no direct synapse from the stimulated cell, and among directly connected pairs, stronger connections do not produce larger responses ^4^. More recent work has shown that signal propagation through indirect pathways contributes substantially to these responses, including between neurons that are directly connected ^5^. However, reproducing these responses required fitting the strengths and signs of synaptic interactions to measured neural activity ^5^. Studies of spontaneous neural activity have also shown that the similarity of neurons’ synaptic input patterns predicts their activity correlations better than direct connection strength does ^6^. Whether the anatomical arrangement of connections is similarly informative about evoked signal propagation, without fitting parameters to neural activity, remains unclear.

If signal propagation depends on this broader network organization, preserving every neuron as a distinct unit may not be necessary. Neurons that participate similarly in those paths may be interchangeable at the level of prediction. *C. elegans* neurons are already grouped into 119 cell classes defined by lineage, morphology and, for some classes, connectivity ^1^. Collapsing the members of a class into a single unit is standard practice in modeling the animal, but whether it preserves what the connectome predicts has not been tested. More generally, this raises the question of how far the connectome can be coarse-grained while preserving its ability to predict signal propagation.

Here we separate these two questions. We first ask how much information about measured signal propagation can be recovered from the anatomical connectome alone, without fitting parameters to neural activity, and which features of the wiring carry that information. We then use this prediction as a functional criterion for coarse-graining, asking which distinctions between neurons can be removed before the connectome loses what it predicts.

At full resolution, signal propagation is predicted by a parameter-free measure of path structure, with nearly all of the improvement over direct connectivity captured within two synaptic steps. Coarse-graining the 302 nodes to the 119 cell classes preserves this prediction despite discarding measurable path structure, whereas size-matched random partitions rapidly degrade it. Connectivity alone identifies many other sufficient partitions, and prediction survives down to roughly nine groups that correspond to recognizable subsystems of the nervous system, although finer structure within these groups remains informative.

This organization is also present across individuals and developmental stages. Partitions inferred from the connectivity of one animal transfer to other animals, even when the particular neuron pairs selected for merging differ. Partitions inferred from newly hatched animals preserve most of the adult connectome’s prediction of signal propagation measured in different adult animals, without using either adult connectivity or neural activity to define them. In the much larger *Drosophila* central brain, lineage-based partitions like-wise preserve path structure substantially better than matched random coarse-graining.

## 2 Results

### Path structure predicts signal propagation

We first asked whether the connectome contains enough information to predict which neuron pairs propagate signal, without fitting any parameters to neural activity. In the brain-wide perturbation atlas of Randi et al. ^4^, 1,098 of 28,989 observed ordered neuron pairs showed a detected response, including 874 pairs with no direct synapse. We scored each pair using a path operator that sums walks through the connectome while progressively down-weighting longer paths (Methods). Predictive performance was quantified by AUC, the probability that a randomly chosen responding pair is assigned a higher score than a randomly chosen non-responding pair, with 0.5 indicating chance performance.

Path structure predicted signal propagation with an AUC of 0.637 across all observed pairs and 0.603 among pairs with no direct synapse (Fig 1B). By comparison, direct synaptic weight reached an AUC of only 0.562 overall and is uninformative by construction among unsynapsed pairs.

**Fig. 1.**
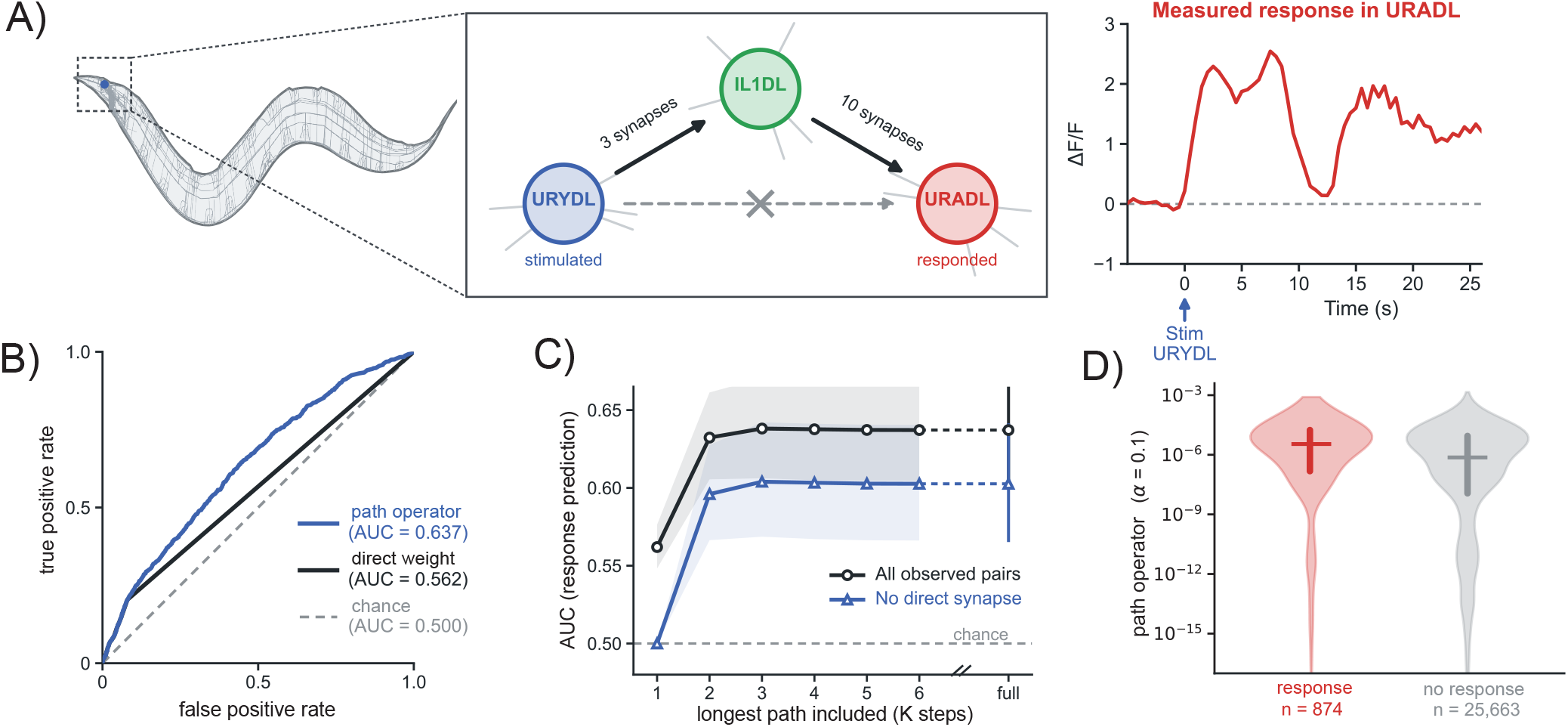
Path structure predicts signal propagation beyond direct synaptic connectivity. **A)** Example of measured signal propagation between two neurons with no direct synaptic connection. Left, the *C. elegans* nervous system in lateral view, with the anterior region containing the example neurons boxed. Center, URYDL and URADL have no synapse in either direction but are linked by a single two-step path through IL1DL, with 3 and 10 synapses along the two edges. Right, URADL calcium response to one stimulation of URYDL, shown as Δ*F/F* relative to the 5 s pre-stimulus baseline. URYDL was stimulated with URADL identified in six events across five recordings; the trace shown is the event in which a response was detected. **B)** Receiver operating characteristic for predicting whether a neuron pair responds, across 28,989 observed ordered pairs, including 1,098 detected responses. The path operator, *R* = (*I* −*αA/ρ*)^−1^ −*I* with *α* = 0.1, reached AUC 0.637, compared with 0.562 for direct synaptic weight and 0.500 at chance. **C)** Predictive performance as progressively longer paths are included using 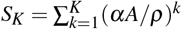. Adding the second step increased AUC by 0.070 across all observed pairs and by 0.096 among pairs with no direct synapse; the third step added only 0.006 and 0.008, respectively. From *K* = 5, the truncated operator matched the full operator to four decimal places. For unsynapsed pairs, *K* = 1 is identically zero and therefore has AUC 0.500 by construction. Bands show 95% cluster-bootstrap intervals from 500 resamples over the 95 stimulated classes. **D)** Path-operator scores for 26,537 observed pairs with no direct synapse, split by whether a response was detected. Bars show the interquartile range crossed at the median. The same separation gives AUC 0.603, 95% CI [0.566, 0.640]. *n* = 874 responding pairs and *n* = 25,663 non-responding pairs.

We next asked whether this advantage could be explained simply by neuronal degree, that is, by how many connections each neuron has. Rewiring the connectome while preserving every neuron’s in- and out-degree reduced performance among unsynapsed pairs from 0.603 to 0.517 ± 0.007 (*z* = 11.4; 10.2 with symmetric gap junctions). Degree-based predictors were similarly weak, reaching an AUC of 0.529 for postsynaptic in-degree and 0.532 for the degree-based expectation of two-hop path count. The predictive information in path structure therefore depends on which neurons are connected through one another, not simply on how many connections they have.

A second concern is that detection depends on how thoroughly each pair was sampled. The observation count, the number of stimulations of the presynaptic neuron that occurred while the postsynaptic neuron was identified, predicted detected responses on its own with an AUC of 0.709, higher than any structural predictor. This sampling is nevertheless nearly uncorrelated with path structure (Spearman *ρ* = − 0.04). Comparing each responding pair only with non-responding pairs of identical observation count lowered the path operator’s AUC only modestly, from 0.637 to 0.615 across all observed pairs and from 0.603 to 0.599 among unsynapsed pairs (Methods). The connectome’s prediction therefore does not arise from its correlation with how often each pair was observed.

Most of this predictive information was already present within two synaptic steps. Among unsynapsed pairs, two-hop path count reached an AUC of 0.601, nearly matching the full path operator at 0.603, and including longer paths produced no measurable improvement (Fig 1C). The connectome’s prediction of signal propagation is concentrated in short paths, particularly those linking each pair through a single intermediate neuron. An example of this is shown in Figure 1A.

Path structure was much less informative about response magnitude. Among the 1,131 detected responses in the released atlas (1,098 of which fall within the observed pairs; Methods), the full path operator correlated only weakly with response magnitude (*ρ* = 0.137), as did two-hop path count (*ρ* = 0.074). Direct synaptic weight is zero for 79.8% of detected responses, so its rank correlation (*ρ* = 0.155) is dominated by a single block of tied values and is not inter-pretable (Supplementary Material). The connectome contains much more information about whether signal propagates than about response strength.

### Cell-class resolution is sufficient for signal propagation

Path structure predicts signal propagation, but that prediction may not require the connectome to be represented at singleneuron resolution. We therefore asked whether it survives coarse-graining. *C. elegans* provides a natural candidate resolution because its neurons are organized into established cell classes, whose members share features such as lineage, morphology and gene expression. Collapsing neurons within these classes reduces the connectome from 302 cells (300 neurons and two muscle cells) to 119 groups (Fig 2A,B).

**Fig. 2.**
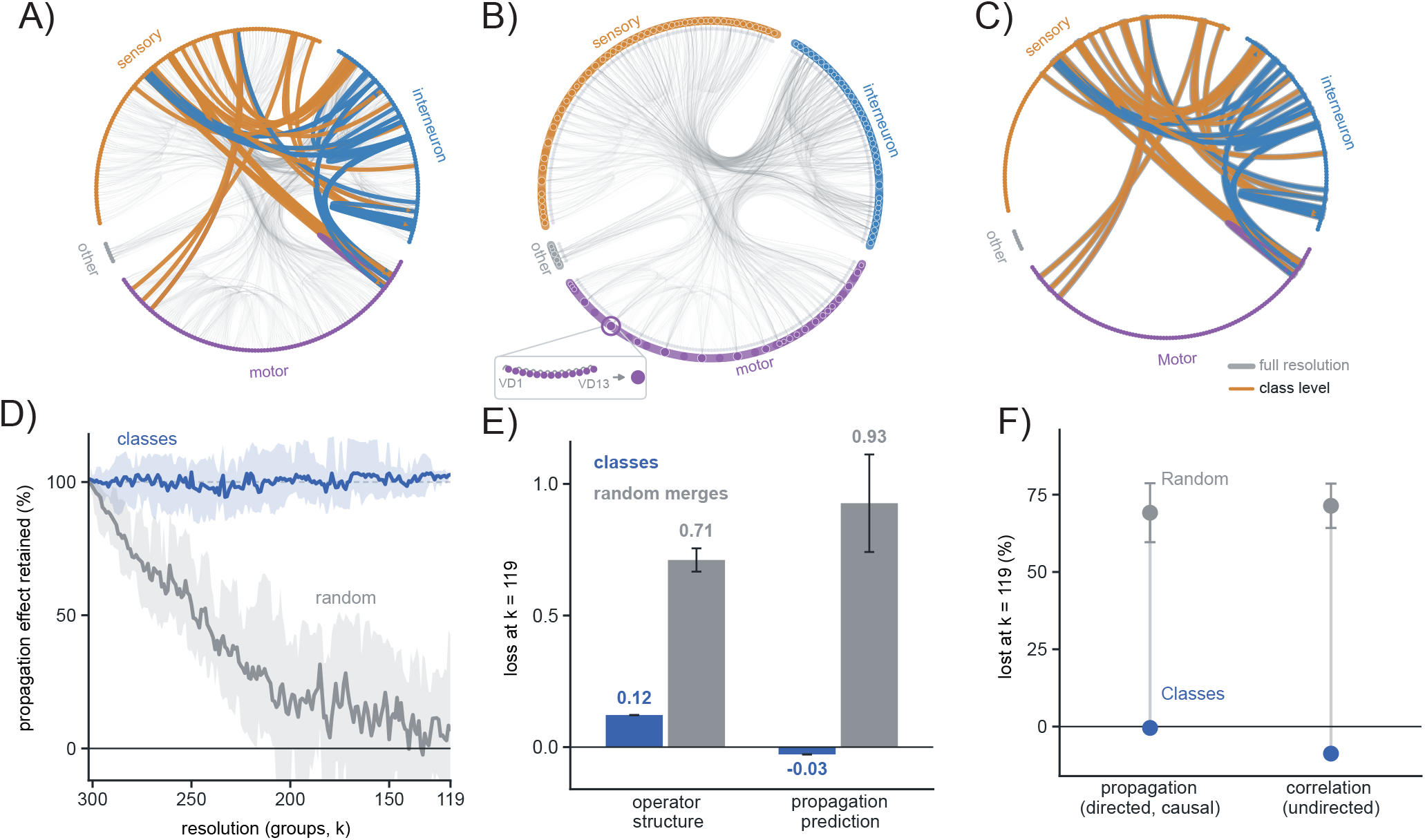
Cell-class resolution preserves the connectome’s prediction of signal propagation. **A)** Full 302-cell connectome, with cells arranged by type. Colored chords show the 70 strongest path-operator predictions between pairs unconnected in either direction, colored by source-cell type. **B)** The same network reduced to 119 biological cell classes, with node size indicating class membership. Inset, the 13 VD neurons merged into one class. **C)** The 70 predictions in **A** scored using the class-level path operator. Gray chords show full-resolution scores; agreement across all 83,970 pairs unconnected in either direction was Spearman *ρ* = 0.80. **D)** Predictive performance during coarse-graining to 119 cell classes compared with size-matched random partitions. Cell classes preserved prediction, whereas random partitions retained only 7% of performance above chance. Shading shows the 2.5–97.5 percentile range over 20 random trajectories. **E)** Structural and predictive loss at *k* = 119. Cell classes had structural loss of 0.12 with essentially no predictive loss, compared with 0.71 and 93% loss of performance above chance for random partitions. Structural loss was evaluated over observed pairs and predictive loss over their unsynapsed subset. Random bars show mean ±s.d. over 20 trajectories. **F)** Cell classes preserved predictions of both signal propagation and activity correlations, whereas random partitions lost 69% and 71% of their respective predictive effects. Loss is normalized to full-resolution performance above chance (AUC−0.5) or full-resolution correlation (*ρ*). Points and error bars show mean ± s.d. over 15 random trajectories.

Neurons within the same cell class show strong similarity in their aggregate connectivity. We quantified this using the equitability defect *γ*, which measures how differently neurons within the same class connect to the rest of the network (Methods). The cell-class partition had a substantially lower equitability defect than size-matched random partitions, which have the same number and sizes of classes but random class membership (*γ* = 0.287 for outgoing and 0.312 for incoming connectivity; *z* = −32.3). Cell classes thus capture a strong regularity in the connectome.

We next coarse-grained the path operator from 302 cells to the 119 cell classes. The resulting class-level operator assigns each neuron pair the value of its two classes and no longer distinguishes between neurons within a class. Despite removing that information, prediction of signal propagation was unchanged. Relative to the full path operator, the change in AUC was +0.001 [95% CI− 0.009, +0.010] across all observed pairs and +0.003 [−0.010, +0.015] among unsynapsed pairs. Expressed as a fraction of the full operator’s performance above chance, cell classes kept 1.00 [0.93, 1.08] of it across all observed pairs and 1.03 [0.91, 1.17] among unsynapsed pairs, so the data exclude losses larger than about 7% and 9% of the effect, respectively. Cell-class resolution is sufficient to preserve the connectome’s prediction of signal propagation (Fig 2D,E).

This preservation depended on which neurons were grouped together. As we progressively coarse-grained the connectome using the biological cell classes, predictive performance changed by less than 0.003, whereas matched random partitions rapidly degraded prediction (Fig 2D). Merging 20% of neurons removed 56% of predictive performance above chance in the random partitions, and by *k* = 180 little remained. Members of the same cell class also evoked more similar sets of responses than neurons from different classes (class-mean Cohen’s *κ*, a chance-corrected agreement, 0.089 compared with 0.014 [−0.002, 0.034] after randomly reassigning class membership, *p* = 0.0005; Supplementary Material). The preservation of prediction also held for a lineage-only partition (*k* = 137) defined without connectivity information (Methods; Supplementary Material), so it does not depend on connectivity having informed the historical class definitions.

### Cell classes preserve prediction while discarding path structure

Preservation of signal-propagation prediction did not mean that coarse-graining preserved the full path structure of the connectome. At *k* = 119, the class-level path operator had a Spearman correlation of 0.88 with the full operator (structural loss 0.12) while producing essentially no loss in predictive performance. Size-matched random partitions had a structural loss of 0.71 and lost most of their predictive performance (Fig 2E).

This separation persisted across resolutions. Starting from the full 302-cell connectome, we progressively merged neurons only within their biological cell classes until each class had been collapsed to a single node. Structural loss increased from 0.067 at *k* = 242 to 0.101 at *k* = 182 and 0.122 at *k* = 119, while the change in AUC remained below 0.003 throughout (Fig 2D).

We next asked how this separation depends on path length. For operators restricted to paths of a single length, the structural advantage of cell classes over size-matched random partitions increased sharply from one to two steps, from 0.27 to 0.54, and changed little thereafter. Averaging one-step connectivity within cell classes already recovered much of the predictive information, improving AUC from 0.562 to 0.608 across all observed pairs and from 0.500 to 0.564 among un-synapsed pairs. Including two-step paths raised performance further, to 0.596 among unsynapsed pairs, while the class-level operator remained effectively lossless relative to the full operator. Two-step paths therefore make the biological organization much more apparent structurally, even though much of the predictive information is already recovered by averaging connectivity within cell classes.

We next asked whether this preservation was specific to the binary measure of signal propagation. The same recordings also provide a continuous measure of pairwise neural activity by correlating the calcium traces of neurons recorded together ^4^. At *k* = 119, the class-level path operator again preserved the relationship between path structure and neural activity, increasing its correlation with measured activity correlations from *ρ* = 0.273 for the full path operator to 0.297, while size-matched random partitions reached only 0.072 ±0.025 (Fig 2F). The result remained after subtracting each neuron’s average response to optogenetic stimulation from its activity trace, showing that the preservation was not driven by neurons sharing the same evoked responses (Methods; Supplementary Material).

We also asked whether sufficiency extended to graded features of the response. The full path operator predicted response amplitude only weakly, and coarse-graining to cell classes left this relationship essentially unchanged, whereas size-matched random partitions removed most of it. The weak prediction of response magnitude is therefore already present at full resolution rather than being introduced by coarse-graining. Response latency could not be tested in the same way because the full path operator carried no reliable information about it.

Across every functional measurement for which the path operator carried detectable information, coarse-graining to cell classes preserved that information while discarding measurable path structure. Cell-class resolution thus preserves the connectome’s functional predictions far more completely than it preserves the underlying network description.

### Connectivity alone identifies sufficient coarse-grainings

Cell-class resolution preserves prediction, but identifying that resolution requires an existing classification of the neurons. We therefore asked whether sufficient coarse-grainings could be identified from connectivity alone.

For every pair of neurons, we calculated how much merging them would increase the equitability defect, which measures the difference in their aggregate connectivity to the rest of the network (Methods). Candidate merges were ranked using connectivity alone, without cell-class labels or measurements of neural activity. Signal propagation was used only afterward to test whether the resulting coarse-grainings preserved prediction.

A complication is that neurons with very few connections can appear similar simply because both have nearly empty connectivity profiles. We therefore restricted the primary analysis to neurons at or above the median total degree. Among these 152 neurons, the 60 disjoint merges ranked best by the criterion produced substantially less predictive loss than size-matched random merges, reducing AUC by 0.0054 compared with 0.0164 (Fig 3C). They also preserved more path structure, with a structural loss of 0.065 compared with 0.175. Part of this structural advantage is expected by construction, since merging neurons with similar connectivity profiles perturbs the path operator less than merging arbitrary neurons. More importantly, these merges also preserve functional prediction and preferentially group neurons from the same biological cell class.

**Fig. 3.**
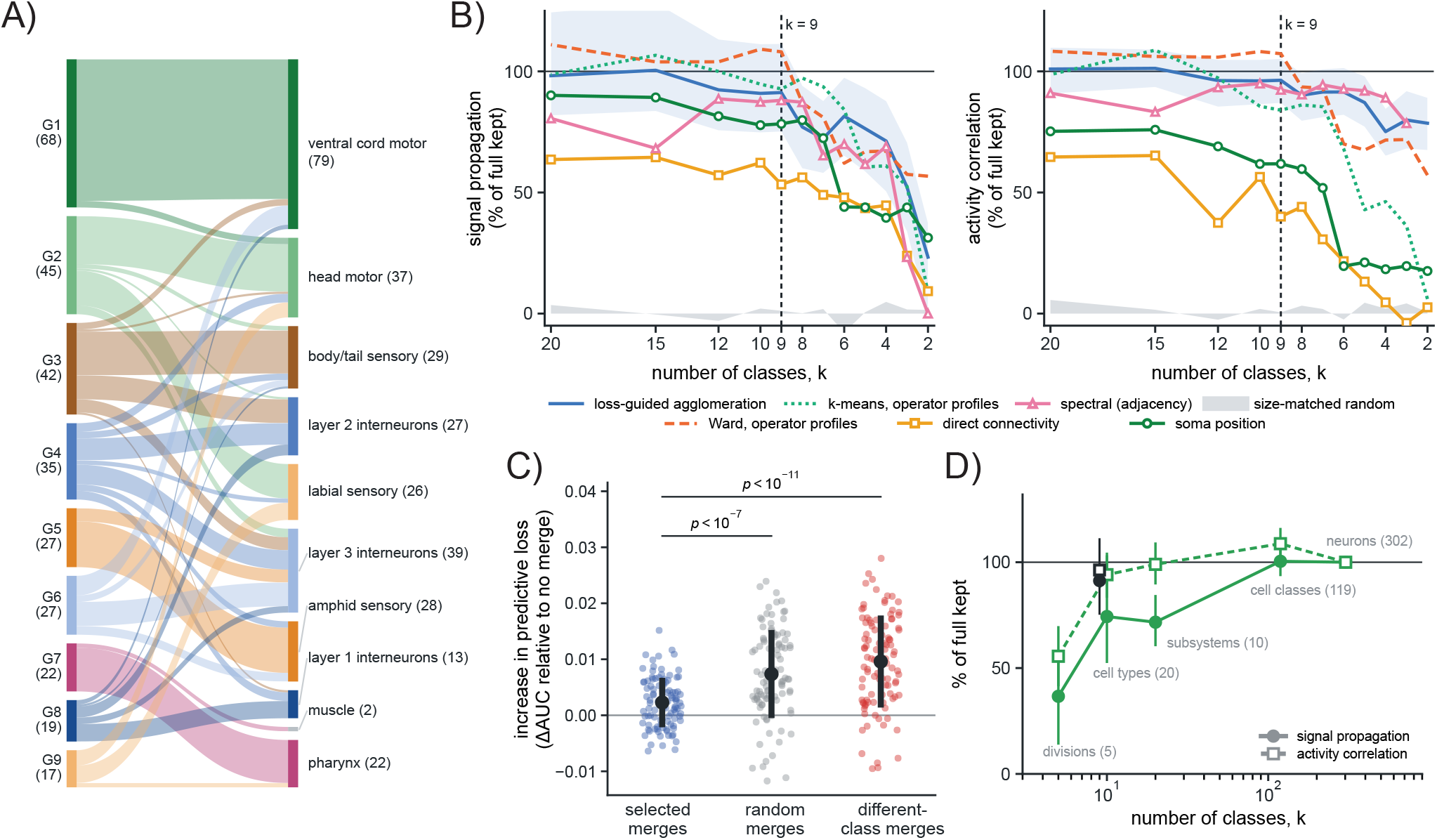
Connectivity identifies coarse-grainings that preserve functional prediction across scales. **A)** The nine groups obtained by loss-guided agglomeration of the path operator, which repeatedly makes the merge that least increases structural loss, compared with canonical biological subsystems. Ribbon width gives the number of shared neurons. The connectivity-defined partition was inferred without biological labels and nevertheless aligned substantially with subsystem identity (adjusted Rand index = 0.49, where 1 is identical partitions; 0.00 for size-matched random partitions). **B)** Predictive performance retained as the connectome is progressively coarse-grained from *k* = 20 to *k* = 2. Partitions derived from path-operator profiles preserved both signal-propagation prediction (left) and activity correlations (right) to approximately *k* = 9, whereas partitions based on direct connectivity, adjacency-spectrum clustering or soma position degraded at finer resolutions. At *k* = 9, loss-guided agglomeration retained 91% of signal-propagation performance and 96% of activity-correlation performance (95% intervals including 100%). Blue shading shows the 95% cluster-bootstrap interval for loss-guided agglomeration. The dashed line marks *k* = 9; gray shading shows size-matched random partitions. **C)** Connectivity alone identifies low-loss merges. Each point shows the increase in predictive loss produced by a set of 30 disjoint merges; connectivity-selected sets were sampled from the 60 best-ranked merges. Black points and bars show mean ±s.d. over the 100 sets. Merges selected from connectivity produced substantially less loss than random or different-class merges (one-sided Mann–Whitney *p <* 10^−7^ and *p <* 10^−11^, respectively). No cell labels or neural activity were used for selection. **D)** Predictive performance retained at successive levels of the canonical biological hierarchy. Cell classes preserved essentially all detectable prediction, whereas cell types, subsystems and broad divisions progressively lost signal-propagation performance (detectably so for cell types and divisions only). The connectivity-defined solution at *k* = 9 (black) retained 91% of signal-propagation and 96% of activity-correlation performance. Bars show 95% cluster-bootstrap intervals.

Although cell-class labels played no role in selecting these pairs, the preferred merges strongly recovered known cell classes. Of the selected pairs, 38% joined neurons from the same cell class, compared with 0.9% expected by chance, a 43-fold enrichment.

We next asked whether the criterion remained effective beyond isolated pairwise merges. We applied it iteratively, recalculating the best merge after each coarse-graining step. Predictive loss remained indistinguishable from zero through 60, 120 and 180 merges, while size-matched random partitions progressively degraded predictive performance. Connectivity alone can therefore identify substantial coarse-graining of the connectome that preserves its prediction of signal propagation.

The resulting connectivity-defined partitions did not perfectly reconstruct the cell classes. As coarse-graining proceeded, they increasingly combined neurons from different cell classes while predictive loss remained indistinguishable from zero. Direct searches for low-loss partitions at *k* = 119 (simulated annealing from cell-class, random and agglom-erative starts; Supplementary Material) reached many structurally different solutions, only weakly overlapping with the biological cell classes (adjusted Rand index 0.12–0.24 across the 14 searches, where 0 is chance agreement and 1 identical partitions), yet almost all preserved signal-propagation prediction (change in AUC among unsynapsed pairs between−0.017 and +0.029 relative to the full operator; a loss was detectable in one of the 14 searches). The cell-class partition is therefore one of many partitions that preserve the connectome’s functional predictions.

### Functional prediction persists at much coarser resolutions

We next asked how far coarse-graining could be taken before prediction was lost. We grouped cells according to the similarity of their path-operator profiles, which describe how strongly each cell reaches, and is reached by, every other cell. We refer to the resulting groups as operator-defined classes. These partitions were constructed from connectivity alone using agglomerative clustering, Ward clustering and *k*-means (Methods).

Across all three methods, no loss of predictive performance for signal propagation or activity correlations was detectable down to approximately nine classes, a 34-fold reduction from the original 302-cell connectome. For the agglomerative solution, nine classes showed no detectable loss for signal propagation or activity correlations, whereas at eight classes loss became detectable for both measures under the primary bootstrap (Fig 3B). Under a two-way bootstrap over stimulated and responding classes, loss first became detectable between eight and five classes depending on the method (Supplementary Material). Structural loss at nine classes was 0.27, compared with 0.96 for size-matched random partitions, while keeping structural loss below 0.05 required at least 235 classes. The resolution at which functional prediction is lost is therefore far coarser than the resolution at which the path operator itself begins to degrade.

Not all methods of coarse-graining preserved prediction equally well. Partitions built from direct connectivity profiles, spectral clustering of the adjacency matrix, or spatial proximity all lost predictive performance at substantially finer resolutions than the operator-defined partitions (Supplementary Material).

### Predictive information is organized hierarchically across the nervous system

The nine operator-defined classes corresponded closely to recognizable subsystems of the nervous system. One contained 20 of the 22 pharyngeal neurons, another 65 of the 68 ventral-cord motor neurons, and a third 21 of the 27 amphid and gas-sensing neurons. To compare this organization with established anatomy, we constructed a nested biological hierarchy from the WormAtlas classification, progressing from neurons to cell classes, cell types, subsystems and broader divisions.

The operator-defined classes agreed strongly with the subsystem partition, with an adjusted Rand index of 0.49 compared with 0.00 for size-matched random partitions (Fig 3A). The subsystem partition itself produced no detectable predictive loss for signal propagation or activity correlations, whereas the five broader divisions lost substantial performance. The estimated loss for signal propagation at sub-system resolution was nevertheless imprecise (0.26 [−0.03, 0.48]). Cell types showed an intermediate pattern, preserving activity correlations but losing some signal-propagation prediction (Fig 3D).

This correspondence was not explained by spatial proximity. Spatial clusters at the same resolution overlapped only weakly with the operator-defined classes, and path-operator structure continued to predict neural activity within spatial clusters. Geometry therefore provides only a partial proxy for the organization that carries predictive information.

We next asked whether this organization appeared under an independent decomposition of the connectome. In the 188 head neurons, operator-defined classes aligned much more strongly with the anatomical communities identified by Dvali et al. than with their signal-propagation communities, and this correspondence remained after excluding the pharynx ^7^. Coarse-graining by the anatomical communities also produced no detectable loss of the path operator’s predictions, whereas size-matched random partitions degraded them substantially (Supplementary Material). Anatomical modular structure can therefore preserve the connectome’s functional predictions even when it differs from the modular organization of measured signaling.

For the nine operator-defined classes, the mean path-operator value between each pair of classes retained essentially all of the full operator’s predictive performance for signal propagation, activity correlations and response amplitude. Neuron-level structure nevertheless remained informative within individual class pairs, where class-level differences are removed by construction. Prediction is therefore dominated by large differences between subsystems, with smaller but real structure remaining within them.

The resolution required therefore depends on the distinctions being predicted. When analysis was restricted to neurons represented across the developmental connectomes, which largely exclude the ventral cord and pharynx, a nineclass partition retained only about half of the path operator’s signal-propagation performance. Predicting differences between major nervous-system subsystems can be done at very coarse resolution, whereas predicting which neurons respond within a subsystem requires finer detail. There is no single sufficient resolution.

### Connectivity-defined organization transfers across animals and development

The preceding analyses identify sufficient coarse-grainings within a single connectome. We next asked whether this organization transfers across animals and across development. The eight developmental connectomes reconstructed by Witvliet et al. span the first larval stage through adulthood, allowing us to test whether connectivity-defined classes found in one animal remain effective in another and how early this organization appears ^8^.

We first asked whether connectivity-defined classes inferred from newly hatched animals could preserve the prediction of signal propagation measured in different, adult animals. We identified these classes independently in each L1 connectome using connectivity alone, then applied the resulting partitions to the adult Cook connectome and tested whether the class-level path operator still predicted signal propagation in the perturbation atlas. The partitions were therefore inferred without access to adult connectivity or neural activity.

At 70 merges among the 161 neurons shared between the developmental and adult connectomes, partitions inferred from L1 animals lost on average only 7.4% of the adult path operator’s predictive performance above chance. Sizematched random partitions lost 21.6%. For comparison, partitions inferred from the adult connectome itself lost 2.4%, while the biological cell classes showed no predictive loss (Fig 4B).

**Fig. 4.**
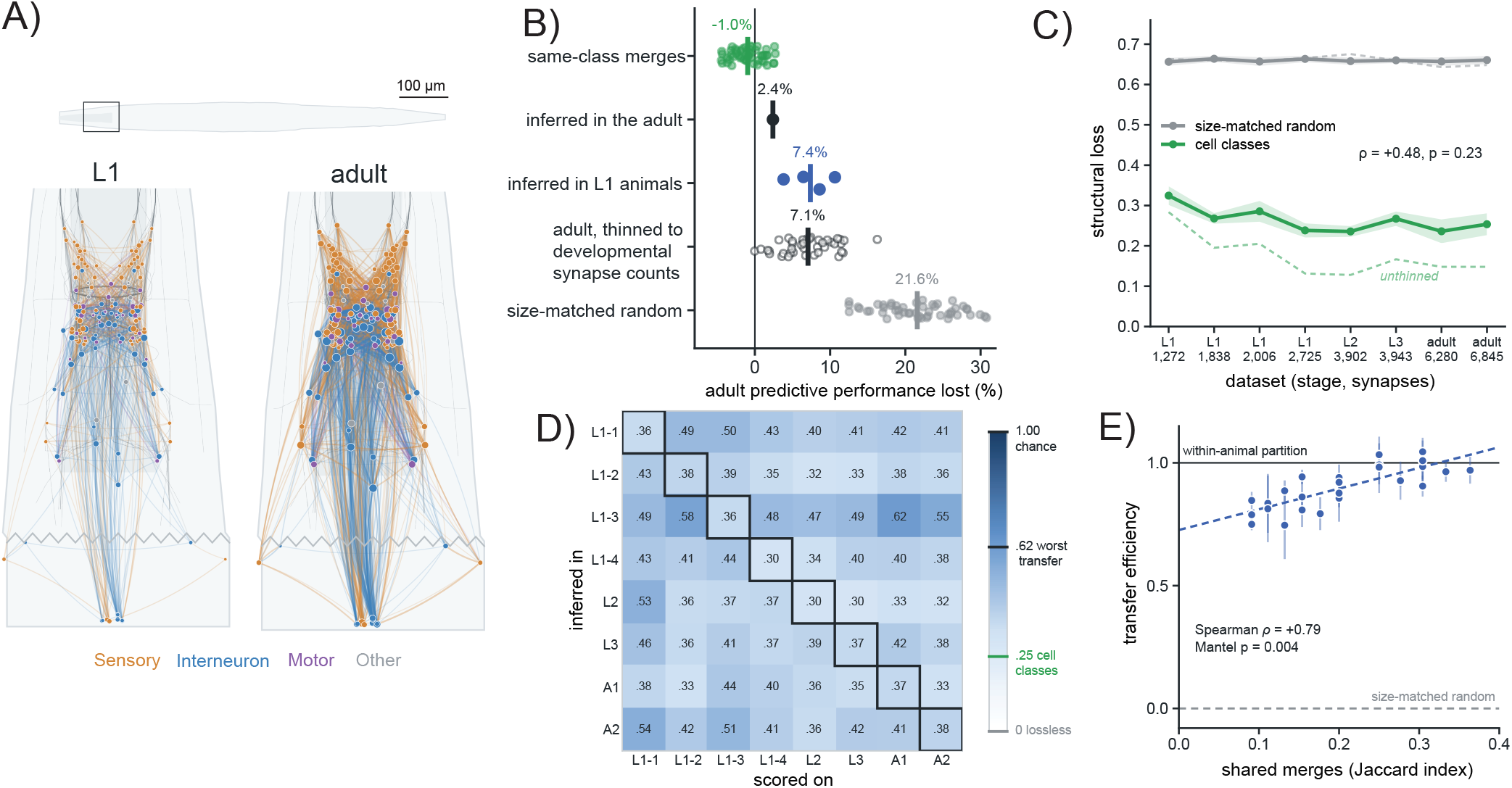
Connectivity-defined organization is present in newly hatched animals and transfers across animals. **A)** L1 and adult developmental connectomes restricted to the 166 cells reconstructed in all eight animals. Nodes are colored by type and connections by presynaptic type; synapse count increases from 1,272 to 6,845. **B)** Loss of adult signal-propagation prediction after applying partitions inferred from different sources. L1-inferred partitions lost 7.4% of predictive performance, close to adults thinned to developmental synapse counts (7.1%) and substantially less than size-matched random partitions (21.6%). Adult-inferred partitions lost 2.4%, while same-class merges showed no loss. Points show individual partitions and vertical bars their mean. **C)** Structural loss of biological cell classes and size-matched random partitions across development after controlling for synapse count. The advantage of cell classes over random partitions showed no detectable developmental trend (Spearman *ρ* = 0.48, *p* = 0.23; *n* = 8 animals). Solid lines and bands show mean ±s.d. after thinning each connectome to 1,200 synapses; dashed lines are unthinned. **D)** Cross-animal transfer of connectivity-defined partitions. Values are structural loss relative to size-matched random partitions on the test connectome; 1 indicates chance and 0 is lossless. Outlined diagonal cells are within-animal partitions; A1 and A2 are the two adults. Transferred partitions retained substantially more structure than random partitions across all animal pairs. **E)** Transfer efficiency increased with overlap in the selected merges (Spearman *ρ* = 0.79, exact Mantel *p* = 0.004), but remained high even when few merges were shared. Across animals, transferred partitions retained 90.8% of the within-animal advantage over random. Each point is one pair of animals, placed at the mean of the two transfer directions, with the vertical line spanning both; the dashed line is a least-squares fit.

At 70 merges, synapse count explained most of the remaining difference between L1- and adult-inferred partitions. The L1 connectomes contain far fewer synaptic contacts than the adult Cook connectome, making the structural criterion less precisely estimated. When the adult connectome was thinned to the same synapse count before inferring the partition, predictive loss increased from 2.4% to 7.1%, nearly matching the 7.4% loss of the L1-inferred partitions (Fig 4B); the same held at 75 merges but not at 40 or 55 (Supplementary Material). At these resolutions, the difference between L1- and adult-inferred partitions is therefore consistent with the sparsity of the available connectomes rather than a different organization of the nervous system.

We then asked whether the structural organization underlying this transfer becomes progressively stronger during development. Synapse count increases from 1,272 to 6,845 across the 166 cells reconstructed in all eight animals (Fig 4A), so raw developmental differences are confounded with network density. We therefore thinned each connectome to a common synapse count before comparing them (Methods).

At every developmental stage, the cell-class partition had a substantially lower equitability defect than size-matched random partitions, lying 15.1 to 18.0 standard deviations below the random expectation, with no detectable trend across development (Spearman *ρ* = 0.071, *p* = 0.87; *n* = 8 animals). Structural loss showed the same pattern, ranging from 0.24 to 0.33 for cell classes compared with approximately 0.66 for size-matched random partitions, with no trend in the difference (*ρ* = 0.48, *p* = 0.23) (Fig 4C). The organization that makes cell-class coarse-graining effective is therefore already present in the earliest connectome, with no detectable strengthening across development after controlling for synapse count.

Its presence at every developmental stage does not imply that the same specific merges are preferred in each animal. We therefore tested transfer directly. For every ordered pair of developmental connectomes, we inferred connectivity-defined classes in one animal, applied that partition unchanged to a second animal, and measured structural loss there. Transfer efficiency, defined as the fraction of the within-animal advantage over size-matched random partitions that was retained after transfer, averaged 90.8% and ranged from 83% to 99%. Mean structural loss increased only from 0.075 within animals to 0.088 after transfer, compared with 0.211 for size-matched random partitions (Fig 4D).

This transfer occurred despite limited agreement in the specific merges selected from different animals. Two animals shared only about one fifth of their selected merges on average (mean Jaccard index = 0.215), even though their partitions transferred with 91% efficiency (Fig 4E). The selected merges were nevertheless strongly aligned with cell-class identity, with 43.8% joining neurons from the same cell class compared with 1.4% expected by chance. Randomly repairing the same selected neurons increased structural loss to 0.208, essentially matching size-matched random partitions at 0.211. Transfer therefore depends on which neurons are paired. Different animals can realize the same coarse-grained organization through different specific merges.

The preferred individual merges vary across animals, but the organization that allows neuron-level distinctions to be discarded is already present in newly hatched animals, with no detectable strengthening across development.

### Lineage-based partitions preserve path structure in *Drosophila*

The *C. elegans* results suggest that biologically organized coarse-graining preferentially preserves path structure. We next asked whether the same principle extends to a much larger nervous system. We analyzed 32,323 neurons of the FlyWire central brain, grouped by hemilineage, a developmental classification of neurons descended from a common progenitor. We then constructed finer partitions by subdividing each hemilineage according to soma position using a preregistered rule (Methods).

Biological partitions preserved substantially more path structure than size-matched random partitions across the full range of resolutions. At the hemilineage level, structural loss was 0.543 compared with 0.997 for matched random partitions, and the advantage persisted at every finer resolution. Because the two species were evaluated on differently sampled cell pairs, we compared them using the ratio of biological to random structural loss. Biological partitions incurred 30–56% of random structural loss across the *Drosophila* hierarchy, compared with 21–25% along the *C. elegans* cell-class trajectory. Preferential preservation therefore extends to a connectome more than two orders of magnitude larger.

Part of this advantage could reflect spatial proximity, because neurons within the same hemilineage often occupy nearby regions of the brain and may therefore share partners. We compared the biological partitions with lineage-blind spatial clusters built from soma position alone at the same resolutions. Spatial clustering recovered 52–70% of the biological advantage across most resolutions in *Drosophila*, but biological partitions consistently preserved more path structure. The same control explained only 30% of the advantage at cell-class resolution in *C. elegans*. The biological advantage therefore generalizes across species, but is more strongly tied to spatial organization in the fly than in the worm.

Unlike in *C. elegans*, the *Drosophila* analysis tests preservation of path structure rather than preservation of functional prediction, because no comparable brain-wide perturbation atlas is available. Biological coarse-graining also remains lossy in both species, with progressively coarser partitions discarding increasing amounts of path structure. The result shared across species is that biologically organized partitions preserve substantially more path structure than matched random coarse-graining at the same resolution.

## 3 Discussion

Direct synaptic connectivity is a poor predictor of measured signal propagation in *C. elegans*, but the connectome becomes substantially more informative once connections are considered in context. Most of this gain was already present within short paths, and averaging connectivity over neurons with similar connectivity profiles recovered much of the same information even before paths were composed. The relevant structural information is therefore contained in how individual synapses are arranged within the surrounding network, in line with the earlier finding that the similarity of neurons’ synaptic input patterns predicts their activity correlations in *C. elegans* better than connection strength or the number of shared inputs ^6^.

Polysynaptic measures are well established in macroscale connectomics ^9;10;11^, where structural network models can predict the spread of activity following stimulation ^12^. Here, polysynaptic structure was likewise informative at the level of individual neurons and synapses. This complements connectome-constrained dynamical models, in which multihop propagation through the anatomical network can account for measured causal interactions once the strengths and signs of direct interactions are learned from neural activity ^5^. The anatomical network alone was much more informative about whether a response occurred than about its magnitude, suggesting that wiring constrains where activity can propagate more strongly than it determines the strength of the resulting response.

Processes that alter effective interactions without changing the underlying wiring may contribute to this separation. Neuromodulation can change the efficacy of anatomically present connections with internal state ^13;14^. However, path structure predicted propagation in *unc-31* mutants as well as in wild type when the strains were compared at equal numbers of stimulations, and the available monoamine and neuropeptide networks did not explain the measured responses. These results do not establish a contribution of dense-core-vesicle signaling to the relationship between anatomical paths and signal propagation. Connectome-constrained dynamical models also account for most causal interactions without adding connections outside the anatomical network ^5^. Extrasynaptic and state-dependent signaling may therefore alter how activity propagates through the anatomical network, although their contribution remains unresolved.

Not every distinction between neurons is equally important for predicting neural activity. Biological cell classes preserved the connectome’s functional predictions despite discarding measurable path structure, whereas matched random coarse-graining removed substantially more of both. The same preferential preservation held for activity correlations and graded response features. What matters is therefore not simply how much structural detail is removed, but which distinctions are removed.

Cell classes were also only one sufficient representation. Connectivity alone identified alternative partitions that preserved prediction, and substantial further coarse-graining remained possible when neurons were grouped according to their path-operator profiles. These coarse partitions aligned with major nervous-system subsystems ^15^, while finer distinctions became necessary when prediction was restricted within the head. The sufficient resolution therefore depends on which distinctions the prediction must preserve rather than being a fixed property of the connectome.

Approximate equitability provides a structural way to formalize this idea ^16;17;18^. In trained recurrent neural networks, connectivity-defined classes recover the organization of task-related dynamics and predict the effects of targeted interventions ^19^. Here, we find that grouping neurons by their connectivity can also preserve predictions of measured neural activity in a biological nervous system. Neurons grouped in this way need not be exactly symmetric or dynamically interchangeable. Their aggregate connectivity need only be similar enough that coarse-graining preserves the network structure relevant to prediction. This approach is related to earlier work on graph fibration symmetries in the *C. elegans* connectome ^20;21^, but does not require exact symmetry. The resulting partitions also overlap with independently identified anatomical communities, even though those communities differ substantially from the communities defined by measured signal propagation ^7^. Anatomical coarse structure can there-fore remain informative about signaling without reproducing the community structure of the signaling network itself.

Sufficient coarse-grainings were not restricted to a single adult connectome. After controlling for synapse count, the cell-class partition was similarly close to equitable from the first larval stage through adulthood ^8^, and partitions inferred from newly hatched animals preserved most of the adult connectome’s predictive performance. Exact preferred merges varied across animals, but the partitions they formed transferred reliably. This is consistent with earlier evidence that a conserved core of connectivity persists alongside variable wiring across individuals ^22^. The distinctions that can be discarded are therefore more stable than the particular synapses or cell pairs through which they are identified in each animal.

The *Drosophila* results suggest that preferential preservation is not unique to *C. elegans*. Lineage-based partitions of the FlyWire central brain ^23;24^ preserved substantially more path structure than matched random partitions across a much wider range of resolutions. Neurons within a hemilineage share developmental origins and often exhibit similar projection patterns and neurotransmitter identities ^25^, providing a biological basis for regularities in their connectivity. Connectome-constrained models have also predicted measured neural responses in the fly ^26;27^, raising the question of whether these predictions could be preserved at lineage-based resolutions. Spatial proximity accounted for more of this advantage in the fly than in the worm, but not all of it. What generalizes across these systems is therefore not a particular sufficient resolution or biological classification, but the preferential preservation of network structure by biologically organized coarse-graining.

Several limitations remain. The static path operator predicted signal propagation and activity correlations better than response magnitude and carried no reliable information about response latency, so the present analysis can identify sufficient resolutions only for functional structure that the operator captures. The perturbation atlas is incomplete and response detection depends on observation count ^4^. It also remains unclear how the relationship between this structural scaffold and neural dynamics changes across internal states.

In *Drosophila*, only structural preservation can currently be tested. More generally, *C. elegans* has an unusually stereo-typed lineage ^3^, so it remains unclear how sharply comparable sufficient resolutions will appear in nervous systems with more variable development.

Connectomes can therefore support prediction at resolutions substantially coarser than the individual neuron. In *C. elegans*, this organization can be recovered from connectivity alone, is already present early in development and transfers across animals despite variation in the underlying wiring. More generally, the useful resolution of a connectome is set not by how much anatomical detail it contains, but by which distinctions in that detail matter for the function being predicted.

## 4 Methods

### 4.1 Connectome data

*C. elegans* analyses used the adult hermaphrodite reconstruction of Cook et al. ^2^, obtained through the OpenWorm Connec-tomeToolbox mirror. Non-neuronal cells were removed by name, leaving 302 nodes (300 neurons and the anal depressor and sphincter muscles) in 119 cell classes (see *Cell classes*). Chemical synapses and gap junctions were combined into a single weighted directed matrix *A*, in which *A*_*i j*_, the connection strength, is the number of synaptic contacts from neuron *i* onto neuron *j*. Rows therefore index presynaptic neurons and columns postsynaptic neurons. Gap junctions are listed in both orientations in this release (all but five of 2,157 entries) and so enter *A* almost exactly symmetrically. The combined matrix contains 29,239 synaptic contacts. The two contact types were also analyzed separately (Supplementary Material).

Developmental analyses used the eight isogenic animals of Witvliet et al. ^8^ from the same mirror, comprising four L1 larvae, one L2, one L3 and two adults. So that the resolution and class-size distribution of every partition were identical at every stage, the node set was fixed at the 166 cells reconstructed in all eight datasets (161 neurons, four CEP sheath glia and one GLR cell; analyses without the glia are in the Supplementary Material). These 166 cells form 71 cell classes under the naming rule below, and their synapse count grows from 1,272 to 6,845 synaptic contacts across the series. Chemical synapses and gap junctions were combined as for the adult, with each gap junction entered in the single orientation recorded in the release.

*Drosophila* analyses used the FlyWire v783 connectome ^23^ with the annotations of Schlegel et al. ^24^. We restricted the analysis to central-brain neurons (super_class = central) with an assigned hemilineage (ito_lee_hemilineage) and a recorded soma position, giving 32,323 neurons. Neurons of the other super-classes were excluded; for the largest of them, optic, sensory and ascending neurons, hemilineage annotations cover 0.2%, 0% and 0%, leaving no connectivity-independent partition to test. *A*_*i j*_ was the synapse count from neuron *i* to neuron *j* in the proofread connection table (54.5 million synapses between 15.1 million connected neuron pairs).

### 4.2 Cell classes

*C. elegans* cell classes were assigned from neuron names. Members of the numbered ventral-cord families (VA, VB, VC, VD, DA, DB, DD and AS) were placed in one cell class by removing the index, so that VA1–VA12 form one cell class. For all other neurons a positional suffix (L, R, DL, DR, VL, VR, D or V) was removed only when the neurons sharing the remaining stem carried a mirror-symmetric set of suffixes (Supplementary Material). The resulting partition agreed exactly with the cell-class assignment distributed with the Randi et al. atlas on all 300 neurons present in both.

Some cell classes were originally defined partly from connectivity ^1^. To exclude this source of circularity, we constructed a partition of lineage-only cell classes, containing only classes definable from lineage and morphology, namely left–right homolog pairs and the serially repeated ventral-cord families, with every other neuron in its own class (*k* = 137).

A canonical anatomical hierarchy was built from the Wor-mAtlas cell type of each neuron, without reference to connectivity, grouping neurons into 19 cell types, nine subsystems and four divisions (sensory, interneuron, motor and pharyngeal; Supplementary Material). The two muscles in the node set form their own class at every level, giving *k* = 20, 10 and 5. The mapping was fixed before any of these partitions was scored. Every multi-member cell class lies within a single cell type and subsystem, so the levels are nested.

### Path operator

Signal can reach a neuron through many paths of different lengths. A path of length *l* from neuron *s* to neuron *r* is any sequence of *l* synaptic steps leading from *s* to *r*, and the entry (*A*^*l*^)_*sr*_ of the *l*-th matrix power sums the product of connection strengths along every such path. Formally these are walks, since a path may revisit a neuron. We summarized path structure with a path operator that adds paths of every length while down-weighting longer ones geometrically,

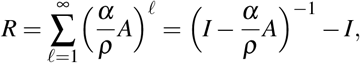

where *ρ* is the spectral radius of *A* (its largest eigenvalue magnitude) and 0 *< α <* 1 sets how rapidly longer paths are discounted. Dividing by *ρ* makes the discount comparable across networks of different size and synapse count and, with *α <* 1, guarantees that the sum converges. Subtracting the identity removes the zero-length path from each neuron to itself. *R*_*sr*_ is the predicted signal propagation from a stimulated neuron *s* to a responding neuron *r*. We used *α* = 0.1 throughout, so that each additional synaptic step multiplies a path’s weight by *α/ρ*; the share of *R*_*sr*_ contributed by each path length still depends on the paths available to that pair. The path operator contains no parameters fitted to neural activity. *R* is the Katz walk-count matrix ^28^, with the discount expressed relative to the spectral radius as in communicability measures ^9^.

To determine how far along the network the predictive information extends, we truncated the sum at path length 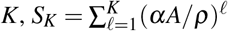, for *K* = 1, …, 6, and compared each truncation with the full path operator. *S*_1_ is a rescaled copy of the direct connection strengths. The *two-step path count* was the number of distinct two-step paths between a pair, (*B*^2^)_*sr*_, where *B* is the binary connection matrix (*B*_*i j*_ = 1 when *A*_*i j*_ *>* 0).

Two controls tested whether prediction depended on the arrangement of connections or only on how many connections each neuron has. First, we scored degree-based predictors directly, including the in-degree of the responding neuron, ∑_*i*_ *B*_*ir*_, and the product of the out-degree of the stimulated neuron and the in-degree of the responding neuron, which is the two-step path count expected in a random network with the same degrees. Second, we recomputed the path operator on 100 degree-preserving surrogates of *A*. Each surrogate was produced by double-edge swaps, each exchanging the targets of two connections (*i* → *j* and *k*→ *l* became *i*→ *l* and *k*→ *j*) while carrying their connection strengths (Supplementary Material). Every neuron therefore retained its exact numbers of incoming and outgoing connections. The two orientations of a gap junction were rewired independently; surrogates that kept them symmetric are described in the Supplementary Material. Unsynapsed pairs were defined from the real connectome in every surrogate, so that identical pairs were scored in every condition.

### 4.4 Measured signal propagation

Signal propagation was taken from the brain-wide perturbation atlas of Randi et al. ^4^, in which single neurons were optogenetically stimulated while calcium activity was recorded across the head. Detected responses were obtained through the atlas API (funconn.princeton.edu; 1,131 wild-type ordered pairs), using the original authors’ detection criterion. Throughout, a *response* means that a pair met this criterion at the number of stimulations it received.

Predicting whether a pair responds requires the pairs that were measured but did not respond, which the released response list omits. We therefore reconstructed the set of observed pairs from the 113 raw wild-type recordings (OSF 10.17605/OSF.IO/E2SYT). For each ordered pair we counted the observation count *O*_*sr*_, the number of stimulations of neuron *s* that occurred in a recording in which neuron *r* was identified, which is the number of opportunities to observe *r* respond to *s*. Neuron labels were retained only when they matched a neuron name in the connectome; glia, annotation notes, unnumbered regions of interest and incomplete identifications were discarded. Pairs with *O*_*sr*_ *>* 0 were considered observed. This gave 28,989 observed ordered pairs, of which 1,098 (3.8%) responded, and matched 97.1% of the released responses (the remaining 33 were dropped). The 26,537 observed pairs with no direct synapse from *s* to *r* (*A*_*sr*_ = 0) were the unsynapsed pairs and contained 874 responses.

Each predictor assigns a single score to every pair, so its predictive performance was measured directly by the area under the receiver operating characteristic curve (AUC), without fitting a model. The AUC is the probability that a randomly chosen responding pair receives a higher score than a randomly chosen non-responding pair, with 0.5 indicating chance. Because observation count strongly affects whether a response is detected, predictors were also scored within strata of identical *O*_*sr*_, comparing each responding pair only with non-responding pairs of the same observation count.

Response magnitude was analyzed only among detected responses, using the Spearman correlation between each predictor and the released response magnitude across the 1,131 detected responses. Response amplitude and latency were measured from the raw recordings (Supplementary Material). Latency was compared with the mean path length implied by the operator, 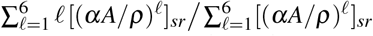, the average number of synaptic steps in the paths from *s* to *r*, each length weighted by its contribution to *R*_*sr*_. At the cell-class partition, predictive loss for each graded observable was expressed as a fraction of the full operator’s correlation and compared with 200 size-matched random partitions. Agreement between the responses evoked by members of a cell class, and the *unc-31* comparison, are described in the Supplementary Material.

### 4.5 Activity correlations

The second dynamical observable was the activity correlation between pairs of neurons, computed as in Randi et al. and Creamer et al. ^4;5^. Within each recording we computed the Pearson correlation between the Δ*F/F* traces of every pair of identified neurons, and averaged these correlations across recordings. Recordings shorter than 100 imaging volumes were excluded, and traces that were entirely missing were dropped from that recording only. We retained the 13,849 unordered neuron pairs recorded together in at least three animals. Because correlation is symmetric, the path operator was symmetrized as *R*_*sr*_ + *R*_*rs*_, and predictive performance was quantified by Spearman’s *ρ* across pairs.

No separate spontaneous recordings are included in the release. Both source studies computed correlations over the full optogenetic recordings, in which a stimulus was delivered approximately every 30 s, so activity correlations and detected responses are computed from the same imaging frames. Shared evoked activity could therefore make the two observables agree. To remove the stimulus-locked component while retaining every frame, we estimated for each neuron in each recording a template spanning 68 volumes (approximately the interval between stimuli). The template was the average over stimulations of the trace following each stimulation onset, expressed relative to its value at onset. This template was subtracted after every onset and correlations were recomputed on the residual traces. Frame excision, sham excision and detrending controls are described in the Supplementary Material.

### 4.6 Equitable partitions and the equitability defect

A partition of *n* neurons into *k* classes was represented by an *n*× *k* indicator matrix *H*, with *H*_*ia*_ = 1 when neuron *i* belongs to class *a* and 0 otherwise. *D* = *H*^⊤^*H* is the diagonal matrix *R* containing the number of neurons in each class. The matrix

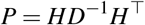

averages within classes. Applied to a vector over neurons, it replaces each neuron’s entry with the mean of its class.

For a connectivity matrix *M*, the product *MP* replaces each neuron’s connections to the members of a class with its average connection to that class, so the *i*-th row of *MP* is neuron *i*’s connectivity profile at the resolution of the partition. A partition is *equitable* when all neurons in a class have the same such profile, that is, when every member of class *a* has the same aggregate connectivity to each class *b* ^16;18^. Exact equitable partitions can be found by color refinement ^29^, and approximately equitable ones by optimization ^30^; the merge criterion and searches below are of the latter kind. We measured departure from exact equitability with the equitability defect

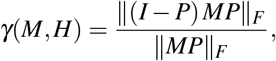

where 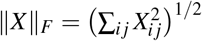 is the Frobenius norm. The numerator measures how far each neuron’s class-level profile deviates from the average profile of its own class, and the denominator scales this by the total connection strength in the profiles. *γ* is zero for an exactly equitable partition and increases as neurons in the same class connect increasingly differently to the rest of the network. Using *M* = *A* compares neurons by their aggregate output (outgoing defect) and *M* = *A*^⊤^ compares them by their aggregate input (incoming defect).

*γ* depends on resolution and class sizes, so it was always compared against size-matched random partitions, generated by randomly permuting class membership across neurons (1,000 partitions for the adult connectome). This defect is the normalized, unsigned counterpart of the equitability defect Γ_signed_ used to identify connectivity-defined classes in trained recurrent networks ^19^. A calibration of *γ* against planted class structure is reported in the Supplementary Material.

### 4.7 Class-level path operator

For an equitable partition, *AH* = *HQ*, with the *k* × *k* quotient matrix

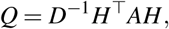

whose entry *Q*_*ab*_ is the average, over neurons in class *a*, of their total connection strength onto class *b. A* then maps vectors that are constant within classes to vectors that are constant within classes, and *Q* reproduces this map exactly; in this sense the network is lumpable under the partition. Because every neuron in a class then sends the same total strength onto each class, the summed path weight from a neuron onto a class is also the same for every member of the neuron’s class, and the quotient reproduces it exactly; what is discarded is how that weight is divided among the members of the receiving class, so individual neuron-pair values of *R* are not reproduced. We coarse-grained the path operator by constructing it on the quotient, *R*_*Q*_ = (−*I αQ/ρ*_*Q*_)^−1^ −*I*, with *ρ*_*Q*_ the spectral radius of *Q*, and assigning each neuron pair the value of its two classes,

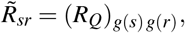

where *g*(*i*) is the class containing neuron *i*. The class-level path operator 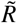 cannot distinguish neurons within a class, since every pair drawn from the same two classes receives the same score. *α* was held at the value used for the full path operator rather than refitted at each resolution, so that any change in prediction reflects the partition rather than re-optimization.

Coarse-graining was evaluated in two ways. *Predictive loss* was the reduction in AUC for predicting signal propagation, 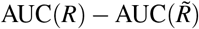, expressed where stated as a fraction of the full path operator’s performance above chance, 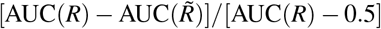. The corresponding fraction for activity correlations was 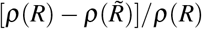, since chance is zero. The two observables were compared only through these fractions, each alongside its own sizematched random partitions, because AUC and Spearman’s *ρ* are not on a common scale. *Structural loss* was 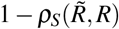, where *ρ*_*S*_ is the Spearman correlation across ordered pairs of distinct neurons. Structural loss is 0 when coarse-graining preserves the rank order of all pair values and approaches 1 when it destroys it. Because the value depends on which pairs are included, structural losses were compared only between partitions scored on the same pairs, which need not be those used for predictive loss. For the adult cell-class partition structural loss was computed over observed pairs, with predictive loss over the unsynapsed subset; structural loss for developmental and connectivity-defined partitions was computed over all ordered pairs.

To determine at which path length cell classes gain their structural advantage, we constructed single-length operators (*A/ρ*)^*l*^ for *l* = 1, …, 4 on the quotient, (*Q/ρ*_*Q*_)^*l*^, assigned each neuron pair the value of its two classes as for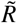, and compared each with the corresponding full operator. The structural advantage at each path length was the mean structural loss of 200 size-matched random partitions minus that of the cell-class partition, on observed pairs (Supplementary Material).

### 4.8 Merge trajectories

To compare cell classes with size-matched random partitions at intermediate resolutions, we constructed matched merge trajectories from *k* = 302 to *k* = 119. In the cell-class trajectory, the 91 multi-member cell classes were merged one at a time in random order, so that *k* fell at each step by one less than the size of the merged class. In the random trajectory, each step merged a class of exactly the same size, drawn at random from neurons not yet merged. At every step the two trajectories therefore had the same resolution and class-size distribution, and differed only in class membership. We generated 20 trajectories of each kind and scored predictive loss on unsynapsed pairs and structural loss at every step. The same trajectories were scored for activity correlations (15 per trajectory type), with signal propagation scored on all observed pairs. At the full cell-class partition, cell classes were additionally compared with 200 size-matched random partitions for each observable.

### 4.9 Connectivity-defined classes

To find sufficient partitions without cell-class labels, we applied the structural criterion pairwise. The merge criterion ranked every possible merge of two neurons by the equitability defect of the partition that places only those two neurons in one class and leaves all others in their own classes. For such a partition *P* differs from the identity only in a 2 × 2 averaging block, and the numerator of the defect reduces to

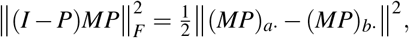

half the squared difference between the connectivity profiles of neurons *a* and *b* after their shared entries are averaged. This closed form, together with a similar one for the denominator ||*MP||_F_*, allowed all 45,451 candidate merges among the 302 cells to be scored exactly. The score for each pair was the mean of its outgoing and incoming defects. Disjoint merges were selected greedily, taking pairs in order of increasing defect and skipping any pair containing a neuron already merged.

Two neurons with very few connections have nearly identical, nearly empty profiles and can appear approximately interchangeable for that reason alone. The primary analysis therefore restricted candidate neurons to those at or above the median total degree (number of distinct presynaptic plus postsynaptic partners; median 29, 152 neurons). Results were also computed with no floor and with floors of 20 and 40 (Supplementary Material). The median floor was chosen after all four had been examined. The 60 disjoint merges ranked best by the merge criterion (*k* = 242) were compared with 30 size-matched random partitions, each consisting of 60 random disjoint pairs drawn from the same candidate neurons, and predictive loss on unsynapsed pairs and structural loss over all ordered pairs were computed for each. For Fig. 3C, 100 sets of 30 merges drawn from the 60 selected were compared with 100 sets of 30 random disjoint pairs and 100 sets of 30 random disjoint pairs of neurons from different cell classes, all from the same candidates (one-sided Mann–Whitney tests). The same-class rate was the proportion of selected merges joining two neurons of the same cell class, with the chance rate given by the probability that two distinct candidate neurons belong to the same cell class.

Beyond approximately 150 merges, disjoint pairs are no longer available, so coarser partitions were built agglomeratively by merging classes. The cost of merging classes *a* and *b*, of sizes *n_a_* and *n_b_*, was

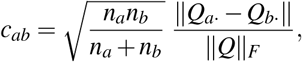

averaged over the outgoing (*Q* computed from *A*) and incoming (*Q* computed from *A*^⊤^) directions. The size weighting reflects that merging two large classes alters the path operator more than merging two small ones, and for two single neurons the cost ranks pairs by the difference in their connectivity profiles. At each step the lowest-cost merge was made and *Q* recomputed. No degree floor was applied. Partitions after 60, 120 and 180 merges were compared with 20 size-matched random partitions.

#### Operator-defined classes

To test how far coarse-graining could extend, classes were built from the similarity of neurons’ path operator profiles by three procedures. In lossguided agglomeration, the agglomerative procedure above was modified so that at each step the 30 lowest-cost merges were each evaluated by the structural loss they produced over all ordered pairs, and the merge producing the lowest structural loss was made. Because this procedure minimizes the structural loss it is later scored on, its structural loss is optimistic by construction; its predictive loss, which it never used, is not. In the other two, each neuron was described by its row and column of the path operator, and these profiles were clustered by Ward hierarchical clustering or by *k*-means (Supplementary Material, which also describes the comparison groupings). Every procedure was run at *k* = 20, 15, 12, 10, 9, 8, 7, 6, 5, 4, 3 and 2, and each partition was scored with the class-level path operator against each observable as a fraction of the full operator’s performance above chance, with 20 size-matched random partitions and cluster-bootstrap intervals (300 resamples). A loss was called detectable when the lower bound of its interval exceeded zero, and the boundary was the largest *k* from which loss in signal propagation or activity correlations, the two observables measured with the narrowest intervals, remained detectable at every coarser resolution. Agreement with the canonical subsystem partition was measured by the adjusted Rand index, the agreement between two partitions corrected for the agreement expected by chance (0 for chance agreement, 1 for identical partitions). *Class-pair means.* To test whether a partition’s classes carry the full operator’s information without the quotient, the full path operator was divided into class-pair means, *R*_block_,whose entry for a neuron pair is the mean of *R* over all ordered pairs of distinct neurons drawn from the same two classes, and *R*_block_ was scored as a predictor in the same way as *R*. Within-class prediction, the comparison with anatomical communities and the search for alternative partitions at *k* = 119 by simulated annealing are described in the Supplementary Material.

### 4.10 Developmental series and synapse count

Synapse count grows more than fivefold across the developmental series, and *γ*, structural loss and the merge criterion all change with synapse count independently of the organization of the connectome. The developmental-series measurements (*γ*, structural loss and merge selection) were therefore repeated after thinning each connectome to a common synapse count of 1,200 synaptic contacts. Thinning drew 1,200 synaptic contacts at random from each connectome with probability proportional to connection strength (a multinomial draw), preserving the relative strengths of connections in expectation. This synapse count lies just below that of the sparsest connectome (1,272 synaptic contacts), so that every animal, including the sparsest, carries comparable sampling noise. Each statistic was averaged over ten thinning replicates. Thinning increases *γ* for every partition, so raw and thinned values were not compared directly. Instead, the cell-class partition was compared at each stage with size-matched random partitions, 100 for the outgoing *γ* (reported as a standardized score) and 20 for structural loss. Trends across the eight datasets were assessed by Spearman correlation with developmental order. The eight datasets are eight different animals, so these trends cannot separate developmental time from individual variation.

### 4.11 Transfer across animals and developmental stages

For structural transfer, the merge criterion was applied independently to each of the eight developmental connectomes. Candidates were restricted to one fixed set of 86 cells at or above the median total degree in the connectivity summed across all eight animals, so that every animal and every comparison drew from the same candidates, and every partition consisted of exactly 30 disjoint pairs (*k* = 136). The connectivity-defined classes found in each animal were then applied unchanged to every other animal and scored by the structural loss they produced there. For each test animal, structural loss was compared across four kinds of partition with identical class-size distributions, namely the withinanimal partition found in that animal itself, the mean of the seven transferred partitions from other animals, 50 partitions of random within-cell-class pairs, and 50 size-matched random partitions drawn from the same candidates. Transfer efficiency was

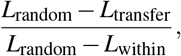

where *L* denotes structural loss, the fraction of the withinanimal partition’s advantage over size-matched random partitions that was kept after transfer, averaged over the eight test animals. Comparisons were made only within a test animal, because structural loss depends on the synapse count of the connectome on which it is scored.

Merge overlap was measured by the Jaccard index (shared merges divided by all distinct merges) between the merge sets of each pair of animals. The chance same-class rate was computed from the 86 candidate cells. To separate which neurons the merge criterion selected from how it paired them, the exact set of neurons selected in each training animal was randomly re-paired (five re-pairings per training animal) and scored on each test animal.

For predictive transfer, connectivity-defined classes found in developmental connectomes were mapped onto the adult Cook connectome by neuron name and scored against measured signal propagation. All 161 neurons shared between the developmental node set and the adult connectome were candidates, and every partition consisted of 70 disjoint pairs (*k* = 232). The degree floor was dropped and the number of pairs chosen after initial runs; every run is reported in the Supplementary Material. Partitions were inferred from each of the four L1 connectomes (averaged), from each of the other developmental connectomes, and from the adult Cook connectome itself. Predictive loss was expressed as a fraction of the full adult path operator’s performance above chance across all observed pairs, and compared with 50 partitions of random within-cell-class pairs and 50 size-matched random partitions. The developmental connectomes have far lower synapse counts than the adult reconstruction, so a less effective partition could reflect synapse count rather than developmental stage. We therefore thinned the adult connections among the candidate neurons to the total synapse count of each developmental connectome (five replicates per synapse count), leaving the remaining adult connections unchanged, inferred partitions from these thinned adult connectomes, and scored them on the complete adult path operator.

### 4.12 *Drosophila* partitions and path operator

Two lineage partitions were defined from annotation alone, the hemilineage partition (*k* = 194) and hemilineages divided by brain side (*k* = 389). Hemilineage subdivisions were generated by a rule fixed before any structural loss was computed (preregistration available in the repository). Within each hemilineage-by-side class of *n* neurons, soma coordinates (standardized across all in-scope neurons) were clustered by Ward hierarchical clustering and the tree was cut into max (1,round(*f n*)) classes. A single parameter *f* was applied to every hemilineage and varied over a fixed grid (*f* = 0.02, 0.05, 0.10, 0.15, 0.20, 0.30, 0.40, 0.55, 0.70, 0.85 and 1.00), giving partitions from *k* = 726 to all 32,323 neurons in their own classes, with reduction factors (*n/k*) from 167 for the hemilineage partition to 1. No alternative clustering method, feature set or cut rule was tried. Hemilineage subdivisions are morphological rather than purely developmental. Cell type, hemibrain type, supertype and cell class annotations were not used because connectivity contributed to their assignment. No brain-wide perturbation atlas exists for the fly, so partitions were evaluated by structural loss only. The full path operator cannot be inverted directly at this scale. Rows of *R* were instead computed by summing the series directly with sparse matrix multiplication up to the first path length *t* with *α^t^*≤10^−6^, which for *α* = 0.1 is length six. *ρ* was estimated by power iteration on |*A|*. Class-level path operators were computed in the same way, normalized by the spectral radius of the full connectome rather than by that of the quotient as in *C. elegans*. The worm and fly implementations were compared by running both on the worm connectome (Supplementary Material). Because out-degree varies widely across neurons, we sampled 200 source neurons stratified by out-degree decile (20 per decile) and 400 random target neurons per source, and scored every partition on the same 80,000 ordered pairs. Each partition was compared with five size-matched random partitions.

Neurons in the same hemilineage tend to lie close together, and nearby neurons can share partners because their arbors occupy the same neuropil. To separate spatial proximity from lineage, we constructed spatial clusters, lineage-blind partitions obtained by mini-batch *k*-means clustering of standardized soma coordinates across all in-scope neurons, at the resolution of every lineage partition and hemilineage subdivision. The share of the biological advantage attributable to spatial proximity was the fraction of the gap in structural loss between size-matched random and biological partitions closed by the spatial clusters. Spatial clusters for *C. elegans* were built in the same way from soma positions in the OpenWorm model, at resolutions along the cell-class merge trajectory. Species were compared through the ratio of biological to size-matched random structural loss, computed over all ordered pairs in *C. elegans* and over the 80,000 sampled pairs in *Drosophila*, because raw structural loss depends on the pairs scored.

### 4.13 Statistics and reproducibility

In the *C. elegans* signal propagation analyses, pairs sharing a stimulated neuron are not independent. Confidence intervals were therefore obtained by a cluster bootstrap that resampled the 95 cell classes of stimulated neurons with replacement, so that the members of a cell class were never treated as independent observations; the stimulated-neuron cell class was thus the unit of replication. Intervals are 95% percentile intervals (200–2,000 resamples; the number is given with each analysis), and paired differences, such as between the full and class-level path operators, were computed within each resample. Intervals that also resample responding-neuron classes or whole recordings are compared in the Supplementary Material, as are the sensitivity of the main results to *α* and to the spectral normalization of the quotient. *Drosophila* intervals were obtained by resampling source neurons.

Partition statistics were always compared with sizematched random partitions, because *γ*, predictive loss and structural loss all change with resolution and class sizes. Standardized scores are *z* = (*x*−*µ*_null_)*/σ*_null_, where *µ*_null_ and *σ*_null_ are the mean and standard deviation across size-matched random partitions or surrogates. The numbers of random partitions, surrogates and trajectories are given with each analysis above. Before interpreting any rank-based or overlap metric computed on sparse binary data, we checked the size of the largest block of tied values, the base rate and how often the metric reached its floor by chance, since such metrics can return confident but uninformative values when most entries are zero.

Analyses were performed in Python using NumPy, SciPy, pandas, scikit-learn and statsmodels, with figures in matplotlib. Code and the preregistered *Drosophila* partition rule are available at https://github.com/cartersale/celegans_connectome_scale.

