## Supplementary Materials for "Connectome structure constrains neural activity across scales"

Supplementary Material

Carter Sale[1,2] and Michael J. Richardson[2]

[1]Modeling and Engineering Risk and Complexity,
Scuola Superiore Meridionale, Naples, Italy

[2]School of Psychological Sciences,
Macquarie University, Sydney, NSW, Australia

### Supplementary Methods

#### Calibration of $\gamma$ and free search

$\gamma$ has no absolute scale, so the cell-class value was calibrated against reference matrices with the same $n = 302$ and the same class-size distribution. In the planted reference, every connection from a neuron in class $a$ to a neuron in class $b$ had the same weight $B_{ab}$, drawn from an exponential distribution with the mean nonzero connection strength of the connectome, so that the cell-class partition is exactly equitable. The noisy planted reference multiplied each entry by $1 + 0.5\,\varepsilon$ with $\varepsilon$ standard normal, giving within-class noise at 50% of the block value. Two structureless references had no class structure: independent standard normal entries, and the connectome with its off-diagonal entries shuffled, which keeps its sparsity and distribution of connection strengths. Each reference was drawn 50 times, and $\gamma$ is reported as the mean of the outgoing and incoming defects. Comparisons across partitions of different resolution were never made, and every $\gamma$ is reported against a null at the same $k$.

Free-search partitions at matched $k$ were obtained by iterative refinement of aggregate-input profiles, minimising $\gamma$ directly, and are reported only as an upper bound on what the criterion itself can achieve.

#### Frame excision, sham excision and detrending

Excision of the volumes surrounding each stimulus is retained as a frame-subsampling check and is not load-bearing for the shared-frames claim, for two measured reasons. The evoked transient does not fit inside the excision window: measured on the stimulated neuron against its own pre-stimulus baseline, $|z|$ runs 2.40 at lag 0 and 7.33 at lag 10, peaks at 13.7 around lags 20–30, and is still 8.2 at lag 79, while stimuli arrive every 67.9 volumes on average. And a frame-matched sham, dropping the same number of windows of the same length at random rather than stimulus-locked onsets, reproduces true excision across three seeds. Sham matching true is consistent both with evoked signal not driving the correlations and with evoked signal being present everywhere, and the transient profile indicates the second is at least partly true.

Losses are never compared across the two observables in raw units. Each is expressed as a fraction of that observable's own headroom above chance, $[\,\text{full} - \text{quotient}\,]/[\,\text{full} - \text{chance}\,]$, and read against a size-matched random arm computed on the same observable, so each observable carries its own calibration. Per-neuron linear detrending was applied as a separate arm, since photobleaching is mild (about 3% over a recording) but shared across neurons and therefore inflates every correlation by a common amount.

#### Functional interchangeability of class members

Agreement between members of a cell class was measured for every pair of stimulated neurons from the same cell class. Responding neurons were restricted to those recorded during stimulation of both members ($O_{sr} > 0$

for each stimulated member $s$), excluding the two stimulated neurons, and comparisons with fewer than 20 such neurons were discarded. Each member's response set was the binary vector of detected responses over these neurons, and agreement was Cohen's $\kappa$, which corrects for the chance agreement expected from the two members' response rates. Because $\kappa$ rises with the number of responses for any pair of neurons, comparisons were repeated after requiring both members to evoke at least three or at least five responses, with the same requirement applied to every comparison. Inference used one value per cell class, the mean $\kappa$ over its member pairs, compared against 2,000 reassignments of class membership among the stimulated neurons with class sizes fixed, and against up to 400 pairs of stimulated neurons from different cell classes (400, 150 and 80 under the three response requirements). Because response vectors are sparse, zero overlap would arise by chance with probability above 0.2 in 78–92% of comparisons, which is why $\kappa$ rather than the Jaccard index was the primary measure. The relation between agreement and structural similarity used, for each cell class, the within-class dispersion of aggregate output relative to size-matched random partitions, with mean observation count and response rate partialled out. Agreement was also scored by the Jaccard index, Spearman correlation and cosine similarity. Class-mean $\kappa$ was 0.089 against 0.014 $[-0.002, 0.034]$ for reassigned classes, and 0.157 and 0.180 against 0.038 and 0.058 under the two response requirements ($p = 0.0005$ for each).

The prediction that more structurally exchangeable classes are more functionally interchangeable was tested by partial Spearman correlation between each class's dispersion ratio and its $\kappa$, controlling exposure and response rate, together with a median split. Minimum detectable effect sizes were computed at $\alpha = .05$ and 80% power for the realised $n$ at each filter.

Every sparse binary metric was first passed through a standing degeneracy check, which reports the largest tie block, the base rate, and how often the metric attains its floor under independence with the observed marginals. Three separate metrics returned confidently wrong answers on sparse binary data before this check was introduced. For a realistically sparse pair, Jaccard's zero-overlap floor is reached 87% of the time by chance, and those uninformative exact zeros are sufficient to reverse the sign of a correlation.

#### Metric parity between the worm and fly analyses

Before comparing the two species we audited every difference between the worm and fly code paths by running both on the worm, where the same matrix passes through each. Spectral radius by dense eigendecomposition against power iteration agreed to $10^{-15}$. The exact dense inverse against a Neumann series truncated at $\alpha^t < 10^{-6}$ differed by at most $1.8 \times 10^{-8}$, with rank correlation 0.999. Dense against sparse quotients agreed to $7 \times 10^{-15}$, and excluding self-pairs moved the statistic by 0.0007.

Only the pair set mattered. Scored on observation-defined pairs the worm's structural loss is 0.1218, and on all ordered pairs 0.1771, a 44% difference, because the observed set contains only pairs whose neurons were both identifiable and imaged together, a biased subset that understates structural loss. Stratified sampling of the kind used in the fly is near-unbiased on the worm, giving 0.1762 against an all-pairs 0.1771, which licenses matching the worm to all pairs for the cross-species comparison while leaving the fly stratified.

The functional and structural losses are therefore quoted together only when both are measured on the observation-defined pairs, where they are 0% and 12%. The all-pairs structural figure of 18% is comparable to the fly and to nothing else, and is never paired with the functional value.

#### Degree floors for the merge criterion

Merges selected by the merge criterion were compared with 60 left–right homolog pairs and with random disjoint pairs, all with the same class-size distribution and evaluated on the same pairs. Two sparsely connected neurons have nearly identical rows, so the analysis was repeated under degree floors at no floor, 20, the median (29) and 40 summed degree, and the median-floor result is reported as primary; this choice was made after results at all four floors had been examined. At the median floor no selected pair contains a below-median-degree neuron. The 40-floor result is not a stronger test. With 83 neurons and 41 available merges the random arm also becomes harmless, falling from +0.0452 to +0.0171, so both arms converge on a floor and the comparison loses contrast before the criterion loses validity.

#### unc-31 comparison

The *unc-31* comparison used the released responses (352 ordered pairs) and 18 raw recordings of *unc-31* mutants from the same study, with observed pairs reconstructed as for wild type (13,185 observed pairs, 343 responses). Wild-type and *unc-31* strains were processed identically, with separate observation denominators built from their own recordings. The reported statistic is the path operator advantage, $\mathrm{AUC}(R) - \mathrm{AUC}(A)$, the difference between the path operator and direct connection strength within strain, and the difference of these advantages across strains, both with intervals from the cluster bootstrap; strains were resampled independently for the difference. To test whether a change was specific to path structure, each predictor's performance above chance ($\mathrm{AUC} - 0.5$) was also compared between strains, as was the ratio of the path operator's performance above chance to that of direct connection strength.

Two controls address the strains' unequal size and detection profile. For sampling, we drew 100 subsamples of 18 wild-type recordings, matching the *unc-31* count, rebuilding the observation denominator from each subsample and recomputing the advantage, and report *unc-31* as a percentile of that distribution. For selectivity, wild-type responses were thresholded to *unc-31*'s detection rate and the advantage recomputed. Extrasynaptic predictors are the monoamine and neuropeptide receptor-expression networks of Bentley et al. [1], scored as pair-level predictors in exactly the same way as the synaptic ones.

Neither control equalizes detection power, because a pair that is stimulated more often has more opportunities to be detected and the wild-type subsets retain response labels detected from all 113 recordings. Two further analyses therefore start from the individual stimulations in the raw recordings of both strains. For each stimulation and each identified responder, the response was the mean fluorescence 1–19.5 s after onset minus the mean over the 5 s before it. The same quantity evaluated at every possible onset of a recording gave each trace a reference distribution, and a stimulation was counted as a response when its deviation from the median of that distribution exceeded the 95th percentile of absolute deviations, so that every trace in both strains has the same exceedance rate by construction. Response magnitude was also expressed in units of the reference distribution's scaled median absolute deviation, in units of the trace's whole-recording standard deviation, as $|\Delta F/F_0|$ with $F_0$ the pre-stimulus mean, and in raw fluorescence units. Each outcome was regressed on the path operator (normal scores of its rank), distance (standardized log distance between somata, or between the nearest points of the two morphologies), the presence of a direct synapse, and the interaction of each with genotype, with a fixed effect per recording. Intervals are from 500 bootstrap resamples of recordings within strain. The model was also fitted to each recording with at least 300 stimulation–responder trials, and the coefficients compared between strains by Mann–Whitney test.

To compare the path operator advantage at equal detection power, detection was recomputed from the raw stimulations. Each stimulation's response was standardized by the median and scaled median absolute deviation of its trace's reference distribution, and a pair's statistic was the sum of these values over its $m$ stimulations divided by $\sqrt{m}$; a pair was detected when the absolute statistic exceeded 2.5 or 3.5. For every pair stimulated in both strains, $m$ was set to the smaller of the two counts and that many stimulations were drawn at random in each strain, 100 times. The advantage was computed in each strain on the same pairs and averaged over draws. Its interval combines one cluster-bootstrap resample of the pairs with each draw, and so covers both the choice of stimulations and the sampling of pairs. This detection statistic is simpler than the one behind the released atlas, with which it agrees at an AUC of 0.815 in wild type.

#### Cell-class naming rule

*C. elegans* cell classes were assigned from neuron names. Members of the numbered ventral-cord families (VA, VB, VC, VD, DA, DB, DD and AS) were placed in one cell class by removing the index, so that `VA1`–`VA12` form one cell class. The numbered pharyngeal neurons `M1`–`M5` and `I1`–`I6` were not merged, because each is a distinct cell class. For all other neurons a positional suffix (L, R, DL, DR, VL, VR, D or V) was removed only when the neurons sharing the remaining stem carried a mirror-symmetric set of suffixes, meaning that every left, right, dorsal-left or ventral-left suffix had its counterpart. This condition prevents spurious cell classes. Greedy suffix removal would, for example, place `AVDL`, `AVDR` and `AVL` under the stem `AV`, whereas `AVL` has no right-hand partner and is its own cell class. Where a name could be parsed in more than one way, the parse giving the largest valid cell class was kept, which assigns all six `RMD` neurons to one cell class rather than splitting off `RMDDL` and `RMDDR`. The resulting partition agreed exactly with the cell-class assignment distributed with the Randi et al. atlas on all 300 neurons present in both.

#### Canonical anatomical hierarchy

A canonical anatomical hierarchy was built from the WormAtlas cell type of each neuron, without reference to connectivity. The 21 WormAtlas types, with the two variants annotated as differing from White et al. folded into their base types, gave 19 cell types. These were grouped into nine subsystems: amphid and gas-sensing neurons; cephalic and labial sensory neurons; body and tail sensory neurons (mechanosensory, touch and phasmid); layer 1, layer 2, and layer 3 with category 4 interneurons; head and sublateral motor neurons; ventral cord and hermaphrodite-specific motor neurons; and the pharynx, including the two neurons linking it to the nerve ring. The subsystems were in turn grouped into four divisions: sensory, interneuron, motor and pharyngeal. The two muscles in the node set form their own class at every level, giving $k = 20$, 10 and 5. The mapping was fixed before any of these partitions was scored. Every multi-member cell class lies within a single cell type and subsystem, so the levels are nested.

#### Degree-preserving surrogates and single-length operators

Each surrogate was produced by ten attempted double-edge swaps per connection, rejecting swaps that would create self-connections or duplicate connections.

For the single-length operators, predictive loss was scored across all observed pairs, because the one-step operator is uninformative among unsynapsed pairs by construction. Intervals on the advantage and on its change between path lengths came from the cluster bootstrap (500 resamples, with 50 of the random partitions rescored in each). Because 94.5% of one-step operator values are zero (91.5% among observed pairs), rank-based structural loss is dominated by tied values at $\ell = 1$, so the increase from one to two steps was also measured as one minus the cosine similarity between class-level and full operator values, which is unaffected by ties.

#### Responses and traces from the raw recordings

Response amplitude and latency were computed from the raw recordings, imaged at 2 volumes $s^{-1}$. For every stimulation, the trace of each other identified neuron from 10 volumes before to 60 volumes after onset was expressed relative to its pre-stimulus mean and divided by the standard deviation of that neuron's full trace, and these traces were averaged over stimulations for each ordered pair. Amplitude was the absolute value of the averaged trace's mean over 1–19.5 s (volumes 2–39) after onset, for the 22,879 observed pairs with at least two stimulations. It predicted detected responses with an AUC of 0.755 and correlated with the released response magnitudes at $\rho = 0.61$ among the 1,025 detected responses with at least two stimulations. Latency was the time at which the averaged trace reached its largest absolute value, for the 1,025 detected responses with at least two stimulations, and was compared with the mean path length implied by the path operator (Methods). Amplitude and latency were likewise scored by Spearman correlation; the operator values have at most 0.8% tied values in either pair set. For response magnitude, the path operator has no tied values among the 1,131 detected responses, and the two-step path count 28%. Direct connection strength is zero for 79.8% of detected responses, so its rank correlation is dominated by a single tied block and was not interpreted.

For activity correlations, traces that were entirely missing (737 region-of-interest-recording combinations, almost all regions annotated as merged and one an identified neuron) were dropped from that recording only. The stimulus template spanned 68 volumes, against a median interval between stimuli of 62 volumes (mean 64).

#### Connectivity-defined classes

To test whether connectivity-defined classes spanning several cell classes were more costly, each multi-member class of the 180-merge partition was labelled pure (all members from one cell class) or mixed and assigned two costs, the predictive loss of merging that class alone and the predictive loss recovered by separating it from the full partition. Because larger classes are both more likely to be mixed and more costly, the primary comparison was restricted to two-neuron classes (one-sided Mann–Whitney test).

In loss-guided agglomeration, every possible merge was evaluated below $k = 20$, where at most 190 merges remain. For Ward clustering and $k$-means, each neuron was described by $\log\bigl(1 + 10^3[R_{i\cdot}, R_{\cdot i}]/\max R\bigr)$, and $k$-means used five runs of ten initializations each. For comparison, classes of the same resolution were built by

$k$-means on direct connectivity profiles, $\log(1 + [A_{i\cdot}, A_{\cdot i}])$, by spectral clustering of the symmetrized adjacency matrix $A + A^\top$, and by $k$-means on soma position. Partitions were scored against signal propagation (all observed pairs and unsynapsed pairs), activity correlations, response amplitude and response magnitude. For $k$-means, spectral and spatial clustering, losses were averaged over the five runs and intervals taken from the first. At $k$ classes the class-level path operator takes at most $k^2$ distinct values, so the largest block of tied values was recorded at every resolution. For loss-guided agglomeration, structural loss was also reported on observed pairs, alongside predictive loss.

Within-class prediction was measured by scoring the full path operator separately within each class pair containing at least 30 observed pairs and averaging these scores weighted by the number of pairs; a predictor that is constant within class pairs scores zero on this measure by construction. Intervals came from the cluster bootstrap (500 resamples for class-pair means, 200 for within-class prediction). Spatial clusters and operator-defined classes were compared by the adjusted Rand index and by scoring each partition's class-pair means within the classes of the other. For the comparison restricted to the nerve ring and head, loss-guided agglomeration (with the 30-merge shortlist at every step) was applied to the adult connectome restricted to the 161 neurons shared with the developmental series, and the class-pair means of the full adult path operator over the resulting nine classes were scored on observed pairs with both neurons in that set.

Community memberships were taken from Tables I and II of Dvali et al. [2], the first hierarchical level of a hierarchical stochastic block model fitted separately to a binarized anatomical network of the head (20 communities) and to the network of detected responses from the same perturbation atlas used here (six communities). Both cover the same 188 head neurons, all of which are in our node set, and the transcribed memberships reproduce the published adjusted Rand index between the two partitions (0.105). The operator-defined classes at $k = 9$, built on the full connectome as above, were restricted to these neurons and compared with each community partition by the adjusted Rand index, against 2,000 random permutations of class membership among the head neurons; for $k$-means, the index was averaged over the five runs. Comparisons were repeated after excluding the 20 neurons of the pharyngeal anatomical community. To score the anatomical communities, head neurons were assigned to their community and every other neuron kept in its own class, and the class-level path operator on this partition was scored against signal propagation, activity correlations and response amplitude on observed pairs with both neurons in the head, as a fraction of the full path operator's performance above chance on the same pairs. Intervals came from the cluster bootstrap (300 resamples), and the comparison was 50 size-matched random partitions that permuted community membership among the head neurons. The signaling communities were not scored in this way, because they were fitted to the responses being predicted.

To test whether sufficient partitions are unique, partitions with $k = 119$ minimizing structural loss over all ordered pairs were sought by simulated annealing, which, unlike agglomeration, can revise earlier assignments. Each proposal either moved one neuron to another class without emptying a class or, in a second search that fixed class sizes to those of the cell classes, exchanged two neurons between classes. Half of the proposals used the class of one of the neuron's 15 most similar neurons in connectivity profile and half the class of a randomly chosen neuron. Proposals were accepted by the Metropolis rule under a temperature decreasing geometrically from $2 \times 10^{-3}$ or $10^{-2}$ to $2 \times 10^{-5}$ over 60,000 proposals, and the best partition found was kept. Fourteen searches were run: in each search space, two from the cell classes and four from random partitions, plus two from the agglomerative partitions at $k = 119$ when class sizes were free. Agreement between partitions was measured by the adjusted Rand index, and predictive loss on unsynapsed pairs by the cluster bootstrap (500 resamples). The search optimizes the structural loss on which it is scored, so it establishes that partitions with lower structural loss exist, not that the best one was found.

### Supplementary Results

#### Signal propagation conditional on observation count

A response is detected after however many stimulations a pair received, so the probability of detection rises with observation count, which alone predicts detected responses at AUC 0.709. Stratifying by quartile does not remove this dependence: the observation count spans 1 to 94 (85 distinct values), and the top quartile alone spans 10 to 94, within which the response rate still rises from 5.6% at 10–14 observations to 34.8% at 50–94. We therefore conditioned on exposure exactly. Observation count was nearly uncorrelated with the path operator (Spearman $\rho = -0.041$ over all observed pairs, $-0.060$ over unsynapsed pairs), so exposure could not account for much of its performance.

Comparing each responding pair only with non-responding pairs of identical observation count lowered the path operator's AUC from 0.637 to 0.615 [0.587, 0.652] across all observed pairs and from 0.603 to 0.599 [0.564, 0.640] among unsynapsed pairs (Table S1). All four structural predictors remained above chance under within-stratum permutation ($p = 0.001$, the smallest attainable with 1,000 permutations), and the ordering of predictors was unchanged. The path operator's advantage over direct connection strength was +0.077 [+0.051, +0.107] within exact strata, the same as without stratification (+0.075). Quartile stratification, by contrast, did not lower the path operator's AUC (0.646), so the earlier quartile control understated how much of the raw performance reflects exposure.

Two models give the same answer. In a logistic model with a separate intercept for every observation count, which assumes no functional form for the effect of exposure, each standard deviation of the path operator's rank multiplied the odds of a detected response by 1.69 [1.52, 1.92] over all pairs and 1.51 [1.34, 1.71] over unsynapsed pairs (likelihood ratio 270 and 139, both $p < 10^{-31}$), and the path operator still added information once direct connection strength was included (likelihood ratio 140, $p = 3 \times 10^{-32}$). In the trial model, in which each stimulation independently evokes a detectable response with probability $q$, the path operator raised the per-stimulation log-odds by 0.509 per standard deviation over all pairs and 0.408 over unsynapsed pairs. Allowing $q$ also to depend on $\log O$ left these nearly unchanged (0.504 [0.403, 0.629] and 0.394 [0.267, 0.510]). The negative coefficient on $\log O$ ($-0.21$ and $-0.27$) indicates that detection rises more slowly with exposure than independent trials would predict, but the path operator's effect does not depend on that assumption. The likelihood-ratio $p$ values treat pairs as independent; the intervals, from the cluster bootstrap over stimulated-neuron cell classes, do not.

Cell classes remained sufficient under exact stratification. The class-level path operator kept 0.97 [0.85, 1.09] of the full operator's performance above chance over all observed pairs and 0.96 [0.78, 1.15] over unsynapsed pairs, against 0.26 and 0.05 for 100 size-matched random partitions ($z = 4.8$ for both), and its difference from the full operator was $-0.003$ [$-0.018$, $+0.010$].

Table S1: **Signal-propagation prediction conditional on observation count.** AUC without stratification, pooled within quartiles of observation count, and pooled within exact observation counts, with 95% cluster-bootstrap intervals (500 resamples) for the last. Observation count scores exactly 0.5 within exact strata by construction.

| | raw | within quartiles | within exact counts |
|---|---|---|---|
| *all observed pairs* (28,989 pairs, 1,098 responses) | | | |
| observation count | 0.709 | 0.577 | 0.500 |
| direct connection strength | 0.562 | 0.561 | 0.538 [0.524, 0.553] |
| two-step path count | 0.627 | 0.630 | 0.594 [0.567, 0.633] |
| path operator | 0.637 | 0.646 | 0.615 [0.587, 0.652] |
| class-level path operator | 0.638 | 0.640 | 0.611 [0.585, 0.646] |
| *unsynapsed pairs* (26,537 pairs, 874 responses) | | | |
| observation count | 0.693 | 0.569 | 0.500 |
| two-step path count | 0.601 | 0.607 | 0.583 [0.549, 0.625] |
| path operator | 0.603 | 0.615 | 0.599 [0.564, 0.640] |
| class-level path operator | 0.605 | 0.612 | 0.595 [0.559, 0.634] |

#### Inference under stronger dependence structures

Observed pairs share stimulated neurons, responding neurons and recordings. The intervals in the main text resample stimulated-neuron cell classes, which addresses the first dependence but not the other two. Table S2 compares that scheme with a two-way bootstrap over stimulated- and responding-neuron cell classes, a bootstrap over recordings, and both together, each with 500 resamples.

Every effect of the path operator was bounded away from zero under every scheme: in none of the 2,000 resamples did its AUC above chance, its AUC within exact observation counts, its advantage over direct connection strength or its correlation with activity correlations reach zero. Two-way resampling widened the intervals by a factor of 1.4–1.8, the largest increase being for activity correlations, where both neurons of a pair are shared with many other pairs. The recording bootstrap produced narrow intervals for signal propagation because the atlas's detection labels, which pool recordings, had to be held fixed; it therefore captures only variation in which pairs were observed and how often, and understates uncertainty for that observable. For activity correlations, which were rebuilt from the resampled recordings, the recording bootstrap gave an interval of [0.205, 0.282] around $\rho = 0.273$. Its downward shift is expected, since fewer distinct recordings contribute to each resample and averaging fewer noisy correlations attenuates their rank correlation with any predictor.

Cell-class sufficiency held under every scheme. The fraction of the full operator's performance kept by cell classes had an interval containing 1 for signal propagation among all and unsynapsed pairs under every scheme, and its excess over size-matched random partitions remained above 0.61 at the lower bound.

The resolution at which operator-defined classes begin to lose predictive performance was less robust than the absence of loss above it. Under every scheme, the loss-guided nine-class partition showed no detectable loss for signal propagation or activity correlations. Under the one-way scheme, loss at eight classes was detectable for both observables (signal propagation kept 0.77 [0.61, 0.92] of full performance). Under the two-way scheme it was not (0.77 [0.49, 1.04]), and loss first became detectable at seven classes for loss-guided agglomeration and $k$-means and at six for Ward clustering, becoming sustained at four to six classes depending on the method (Table S3). The statement supported under all schemes is therefore that operator-defined partitions show no detectable loss down to nine classes; the point at which loss becomes detectable lies between eight and about five classes depending on how dependence among pairs is modelled.

Table S2: **95% intervals under four resampling schemes.** AUC and $\rho$ are shown as performance above chance. Fractions are the class-level path operator's performance relative to the full operator's.

| | estimate | source | source × target | recording | recording + two-way |
|---|---|---|---|---|---|
| path operator, AUC − 0.5, all pairs | 0.137 | [0.103, 0.172] | [0.084, 0.187] | [0.136, 0.144] | [0.090, 0.186] |
| unsynapsed pairs | 0.103 | [0.065, 0.141] | [0.049, 0.151] | [0.099, 0.110] | [0.053, 0.154] |
| within exact observation counts | 0.115 | [0.085, 0.154] | [0.050, 0.164] | [0.107, 0.135] | [0.064, 0.176] |
| path operator − direct connection strength | 0.075 | [0.048, 0.103] | [0.037, 0.111] | [0.072, 0.081] | [0.040, 0.112] |
| median-score ratio, unsynapsed (Fig. 1D) | 4.7 | [2.9, 8.2] | [2.2, 11.6] | [4.5, 5.2] | [2.1, 12.4] |
| path operator, $\rho$ with activity correlations | 0.273 | [0.234, 0.316] | [0.206, 0.351] | [0.205, 0.282] | [0.179, 0.339] |
| cell classes, signal propagation | 1.00 | [0.93, 1.08] | [0.86, 1.17] | [0.97, 1.02] | [0.86, 1.16] |
| unsynapsed pairs | 1.03 | [0.91, 1.17] | [0.76, 1.41] | [0.98, 1.06] | [0.78, 1.34] |
| activity correlations | 1.09 | [1.02, 1.17] | [1.01, 1.24] | [1.04, 1.12] | [0.98, 1.26] |
| minus random, unsynapsed | 0.89 | [0.75, 1.09] | [0.62, 1.39] | [0.85, 0.94] | [0.61, 1.36] |
| loss-guided $k = 9$, signal propagation | 0.91 | [0.74, 1.10] | [0.66, 1.22] | [0.89, 0.94] | [0.63, 1.23] |
| activity correlations | 0.96 | [0.86, 1.07] | [0.81, 1.15] | [0.91, 1.01] | [0.81, 1.15] |
| loss-guided $k = 8$, signal propagation | 0.77 | [0.61, 0.92] | [0.49, 1.04] | [0.75, 0.81] | [0.47, 1.05] |
| activity correlations | 0.90 | [0.80, 0.99] | [0.73, 1.07] | [0.82, 0.96] | [0.72, 1.09] |

#### Sensitivity to the path discount $\alpha$

The path operator contains one fixed hyperparameter, $\alpha$, which sets how strongly longer paths are discounted. It is not fitted to activity, but the class-level path operator is computed at $\alpha$ and the operator-defined classes cluster profiles of the operator, so we rebuilt every $\alpha$-dependent result from scratch at seven values spanning two orders of magnitude ($\alpha = 0.01$ to 0.9; Table S4). At each value, the loss-guided agglomeration, Ward clustering and $k$-means (three initializations) were rerun on the operator at that $\alpha$, and every partition was

Table S3: **Loss of operator-defined partitions under two-way resampling.** Loss as a fraction of the full path operator's performance above chance, with 95% two-way bootstrap intervals (500 resamples). * Lower bound above zero. $k$-means from one initialization.

| | loss-guided | | Ward on operator | | $k$-means on operator | |
|---|---|---|---|---|---|---|
| $k$ | propagation | correlation | propagation | correlation | propagation | correlation |
| 10 | 0.09 | 0.04 | −0.09 | −0.08 | −0.04 | −0.09 |
| 9 | 0.09 | 0.04 | −0.08 | −0.07 | −0.05 | 0.08 |
| 8 | 0.23 | 0.10 | 0.13 | 0.06 | 0.03 | 0.29 |
| 7 | 0.28* | 0.09 | 0.19 | 0.07 | 0.24 | 0.42* |
| 6 | 0.19 | 0.09 | 0.38* | 0.30* | 0.23 | 0.27 |
| 5 | 0.24 | 0.13 | 0.33 | 0.32* | 0.42* | 0.59* |
| 4 | 0.29 | 0.25* | 0.33 | 0.28* | 0.49* | 0.67* |
| 3 | 0.48* | 0.20 | 0.43* | 0.28* | 0.50* | 0.68* |

scored with the class-level path operator at the same $\alpha$, using the boundary rule of the main text with 200 cluster-bootstrap resamples. The two-step path count, which does not depend on $\alpha$, reached AUC 0.627.

The full operator's prediction changed little over most of the range and declined only as $\alpha$ approached 1. Signal-propagation AUC was 0.640 at $\alpha = 0.01$, 0.637 at the value used throughout, and 0.630 at 0.3, falling to 0.591 at 0.9; its correlation with activity correlations was 0.27 from 0.01 to 0.7 and 0.24 at 0.9. The value used in the paper was therefore not selected to maximize prediction; smaller values predicted marginally better.

Cell-class sufficiency did not depend on $\alpha$. At every value, the class-level path operator on cell classes lost at most 0.01 of the full operator's performance for signal propagation and none for activity correlations, while size-matched random partitions lost 0.66–0.73 and 0.73–0.75.

The operator-defined boundary was stable from $\alpha = 0.01$ to 0.7. Across the 18 combinations of method and $\alpha$ in this range, sustained loss began at 10 or fewer classes in every case and at 8 or fewer in 15 of them (in one, none was detectable down to four classes). Four combinations showed detectable loss already at nine classes in one observable (loss-guided agglomeration at $\alpha = 0.5$ for signal propagation; Ward clustering at 0.3 and 0.7 and $k$-means at 0.7 for activity correlations). At $\alpha = 0.9$, where long paths dominate the operator, all three methods lost performance by 15–20 classes. The operator-defined partitions themselves changed with $\alpha$: the nine-class partition at each value agreed with the one at $\alpha = 0.1$ with an adjusted Rand index of 0.28–0.64 across methods, and with the canonical subsystems at 0.24–0.49. Coarse sufficiency near nine classes therefore does not depend on the particular value of $\alpha$ below about 0.7, although which nine-class partition is found does, consistent with the non-uniqueness of sufficient partitions reported in the main text.

Table S4: **Sensitivity to $\alpha$.** Full path operator performance; loss of the class-level operator on cell classes as a fraction of full performance (negative values are gains); the resolution from which loss in signal propagation or activity correlations is detectable at every coarser resolution scored (–, none down to $k = 4$); and agreement of the loss-guided nine-class partition with that at $\alpha = 0.1$ and with the canonical subsystems.

| | full operator | | | cell-class loss | | sustained loss from $k$ | | | loss-guided $k = 9$, ARI | |
|---|---|---|---|---|---|---|---|---|---|---|
| $\alpha$ | AUC | AUC unsyn. | $\rho$ corr. | propagation | correlation | loss-guided | Ward | $k$-means | vs $\alpha = 0.1$ | vs subsystems |
| 0.01 | 0.640 | 0.607 | 0.271 | 0.00 | −0.09 | – | 7 | 7 | 0.46 | 0.27 |
| 0.03 | 0.640 | 0.606 | 0.272 | 0.00 | −0.09 | 4 | 6 | 8 | 0.45 | 0.36 |
| 0.1 | 0.637 | 0.603 | 0.273 | 0.00 | −0.09 | 8 | 7 | 8 | 1 | 0.49 |
| 0.3 | 0.630 | 0.593 | 0.274 | −0.01 | −0.08 | 8 | 10 | 8 | 0.59 | 0.37 |
| 0.5 | 0.622 | 0.584 | 0.272 | −0.01 | −0.08 | 10 | 6 | 7 | 0.55 | 0.47 |
| 0.7 | 0.611 | 0.572 | 0.266 | 0.00 | −0.06 | 7 | 4 | 10 | 0.42 | 0.38 |
| 0.9 | 0.591 | 0.554 | 0.243 | 0.01 | −0.02 | 20 | 15 | 20 | 0.39 | 0.29 |

#### Spectral normalization of the class-level path operator

In *C. elegans* the class-level path operator normalizes the quotient $Q$ by its own spectral radius $\rho_Q$; in *Drosophila* it is normalized by the spectral radius $\rho$ of the full connectome. The two coincide for an equitable partition, because $AH = HQ$ then implies that the leading eigenvalue of $A$ is an eigenvalue of $Q$, so they can differ only to the extent that a partition departs from equitability. We measured that departure and repeated the key results in each species under the other species' convention.

In the worm, the partitions carrying the main results are close to spectrally exact: $\rho_Q/\rho$ was 0.978 for cell classes, 0.979 for lineage-only cell classes and 0.997 for the 60 merges selected by the merge criterion, against 0.70, 0.72 and 0.90 for their size-matched random partitions (Table S5). Operator-defined classes at nine classes gave 0.85–0.88, against 0.56 for random partitions. Holding the normalization at the full connectome's $\rho$ left every biological result unchanged to the third decimal place. The class-level operator on cell classes kept 1.005 of the full operator's signal-propagation performance (1.004 with $\rho_Q$), 1.029 among unsynapsed pairs (1.028) and 1.087 for activity correlations (1.087), with structural loss 0.177 over all ordered pairs in both. Random partitions were also barely affected. Their effective discount per step falls from 0.1 to about 0.07 under the fixed normalization, too small a change to reorder many pairs, and they retained 0.29 and 0.30 of signal-propagation performance under the two normalizations. The advantage of the biological partitions over random partitions was therefore essentially the same under both conventions: for cell classes, $z$ was 5.2–9.4 for signal propagation, activity correlations and response amplitude under either normalization, and structural loss lay 13–18 standard deviations below random under both (Table S5).

The operator-defined boundary was also unchanged in its main feature. Under the fixed normalization no method showed detectable loss at nine classes, Ward clustering and $k$-means lost performance from the same resolutions as before (seven and eight classes), and loss-guided agglomeration, rebuilt with the fixed normalization in its objective, from five rather than eight classes, within the range reported in the main text.

Table S5: ***C. elegans* results under the two normalizations.** Fraction of the full path operator's performance kept by the class-level operator normalized by its own spectral radius ($\rho_Q$) or by the full connectome's ($\rho$), with the mean over 100 size-matched random partitions (random disjoint pairs from the same neurons for the merge criterion). Structural loss is over all ordered pairs.

| | cell classes | | lineage-only | | merge criterion (60) | |
|---|---|---|---|---|---|---|
| | $\rho_Q$ | $\rho$ | $\rho_Q$ | $\rho$ | $\rho_Q$ | $\rho$ |
| $\rho_Q/\rho$ (random) | 0.978 (0.70) | | 0.979 (0.72) | | 0.997 (0.90) | |
| signal propagation | 1.004 | 1.005 | 1.069 | 1.069 | 0.963 | 0.963 |
| random | 0.302 | 0.292 | 0.335 | 0.343 | 0.882 | 0.887 |
| unsynapsed pairs | 1.028 | 1.029 | 1.143 | 1.144 | 0.948 | 0.948 |
| random | 0.061 | 0.045 | 0.064 | 0.073 | 0.839 | 0.849 |
| activity correlations | 1.087 | 1.087 | 1.051 | 1.051 | 0.958 | 0.958 |
| random | 0.270 | 0.243 | 0.306 | 0.295 | 0.884 | 0.888 |
| response amplitude | 0.998 | 0.998 | 1.133 | 1.134 | 0.974 | 0.974 |
| random | 0.179 | 0.197 | 0.258 | 0.251 | 0.902 | 0.908 |
| structural loss | 0.177 | 0.177 | 0.159 | 0.159 | 0.065 | 0.065 |
| random | 0.755 | 0.761 | 0.704 | 0.705 | 0.176 | 0.177 |

In the fly the lineage partitions are much further from spectral exactness: $\rho_Q/\rho$ was 0.45 for hemilineages and 0.46 for hemilineages divided by side, rising to 0.56, 0.66 and 0.88 for hemilineage subdivisions at $f = 0.05$, 0.2 and 0.55, against 0.21, 0.21, 0.23, 0.30 and 0.57 for size-matched random partitions. Normalizing every fly quotient by its own spectral radius, as in the worm, nevertheless changed structural loss by at most 0.005 for any lineage partition and 0.004 for its random partitions, across all 13 resolutions from hemilineages to single neurons (Table S6). This insensitivity is consistent with the operator at $\alpha = 0.1$ being dominated by paths of one and two steps, whose rank order a moderate rescaling of the per-step discount changes little. Neither species' conclusions depend on which normalization is used.

#### Path operator advantage in *unc-31* mutants

The *unc-31* recordings contain far fewer observations than wild type (Table S7): 18 rather than 113 recordings, 13,185 rather than 28,989 observed pairs, and a mean observation count of 2.8 rather than 7.4 per pair. Within each strain, the path operator outperformed direct connection strength, but the advantage fell from +0.075 [+0.048, +0.103] in wild type to +0.033 [+0.002, +0.062] in *unc-31* mutants. The difference between strains was +0.042 [+0.002, +0.083] (one-sided $p = 0.021$, two-sided $p = 0.041$).

Two explanations unrelated to dense-core-vesicle release were first tested. The first is sampling. Wild-type observed pairs rebuilt from 100 random subsets of 18 recordings, which gave a mean observation count of

Table S6: ***Drosophila* structural loss under the two normalizations.** Structural loss of each lineage partition and the mean of five size-matched random partitions, with each quotient normalized by the full connectome's spectral radius $\rho$ (as in the main text) or by its own $\rho_Q$.

| | | lineage partition | | random | |
|---|---|---|---|---|---|
| partition | $k$ | $\rho$ | $\rho_Q$ | $\rho$ | $\rho_Q$ |
| hemilineage | 194 | 0.543 | 0.548 | 0.997 | 0.997 |
| hemilineage × side | 389 | 0.556 | 0.559 | 0.991 | 0.991 |
| subdivisions, $f = 0.02$ | 726 | 0.483 | 0.481 | 0.975 | 0.974 |
| $f = 0.05$ | 1,663 | 0.439 | 0.439 | 0.971 | 0.969 |
| $f = 0.10$ | 3,261 | 0.412 | 0.411 | 0.942 | 0.941 |
| $f = 0.20$ | 6,478 | 0.368 | 0.366 | 0.866 | 0.865 |
| $f = 0.40$ | 12,935 | 0.295 | 0.293 | 0.742 | 0.739 |
| $f = 0.55$ | 17,785 | 0.235 | 0.234 | 0.629 | 0.625 |
| $f = 0.85$ | 27,477 | 0.098 | 0.097 | 0.324 | 0.323 |

2.3, below that of the mutant, kept an advantage of +0.079 [+0.068, +0.088]; the mutant value lay below every subsample. The second is detection. Detected *unc-31* responses were larger than wild-type responses (median magnitude 25 against 14; Mann–Whitney $p = 3.5 \times 10^{-21}$), as expected if detection in the mutant is more selective, but restricting wild-type responses to the largest 69%, matching the mutant response rate, left the advantage at +0.075. Neither control moved the advantage by more than 0.005, against a difference of 0.042 between strains.

The reduction was not specific to path structure. Every connectome-based predictor lost about half its performance above chance: direct connection strength from 0.062 to 0.032, the path operator from 0.137 to 0.064, and the path operator among unsynapsed pairs from 0.103 to 0.042. The ratio of path-operator to direct-connection performance above chance was therefore almost unchanged, 2.21 in wild type and 2.04 in the mutant. The monoamine and neuropeptide networks predicted responses at chance in both strains (AUC 0.496–0.503). The smaller absolute advantage in the mutant therefore reflects a proportional loss of the relationship between connectivity and response, rather than a selective loss of the contribution of multi-step paths.

These comparisons establish that the released detections of the two strains differ, but not that responses do. A pair that is stimulated more often has more opportunities to be detected, and neither control removes that dependence: the recording subsets changed which pairs counted as observed while keeping response labels detected from all 113 recordings. The released detections themselves show its size. Among pairs stimulated at most five times, the response rate was 2.0% in wild type (16,615 pairs) and 2.4% in the mutant (11,772 pairs). The wild-type excess arose entirely among pairs stimulated more than five times, 6.2% against 4.0%, which make up 43% of wild-type observed pairs and 11% of mutant ones. The stronger test is whether the strains differ when each stimulation is counted once.

They differed little (Table S8). The raw recordings gave 213,353 stimulation–responder trials in wild type and 36,201 in the mutant, from recordings of similar noise and fluorescence level. A stimulation evoked a response in 9.8% of trials in wild type and 8.7% in the mutant when the pair was directly connected, and the two strains converged as the shortest path between the pair lengthened, reaching 3.9% and 4.2% at four or more steps, against 5% expected without any response. In the trial-level model, no interaction between genotype and the path operator was resolved for response occurrence, whether distance was measured between somata (+0.002 [−0.003, +0.007] per standard deviation of path-operator rank) or between morphologies (+0.000 [−0.005, +0.005]), and none was resolved for a direct synapse. Occurrence fell slightly more steeply with distance between morphologies in wild type (−0.004 [−0.008, −0.001]), but not with distance between somata (−0.001 [−0.008, +0.004]). For response magnitude the result depended on the scale. The interaction with the path operator was positive in units of the whole-recording standard deviation (+0.019 [+0.005, +0.031]) and as $|\Delta F/F_0|$ (+0.006 [+0.002, +0.010]) when distance was measured between somata, but was not resolved for either with distance between morphologies, nor in noise units or raw fluorescence under either distance. Coefficients fitted to single recordings did not differ between strains for any term (Mann–Whitney $p \geq 0.33$; 98 wild-type and 13 mutant recordings).

The difference in path operator advantage did not survive equal detection power either. For the 12,520 pairs stimulated in both strains, detection was recomputed from the same number of stimulations in each (2.6

per pair on average). Detection rates were then equal, 4.7% in each strain at the stricter threshold, and the path operator outperformed direct connection strength in both: by +0.032 [+0.004, +0.058] in wild type and +0.043 [+0.014, +0.078] in the mutant, a difference of −0.011 [−0.046, +0.025]. At the looser threshold the difference was +0.005 [−0.019, +0.028]. This detection statistic is cruder than that of the released atlas; with all stimulations it gave a wild-type advantage of +0.031 to +0.035 rather than +0.075, so the test establishes the absence of a strain difference under a detector applied identically to both strains, not what the released detection would give on matched stimulations.

The smaller advantage in the released *unc-31* detections therefore reflects how often each pair was stimulated rather than the loss of dense-core-vesicle release. Counted per stimulation, responses followed path structure in both strains, and these data do not resolve a dependence of that relationship on genotype.

Table S7: **Signal-propagation prediction in wild type and *unc-31* mutants.** AUC over all observed pairs unless stated. Intervals are 95% cluster-bootstrap intervals.

| | wild type | *unc-31* |
|---|---|---|
| recordings | 113 | 18 |
| observed pairs | 28,989 | 13,185 |
| responses (rate) | 1,098 (3.8%) | 343 (2.6%) |
| mean observation count | 7.36 | 2.75 |
| direct connection strength | 0.562 | 0.532 |
| path operator | 0.637 | 0.564 |
| path operator, unsynapsed pairs | 0.603 | 0.542 |
| monoamine network | 0.501 | 0.496 |
| neuropeptide network | 0.503 | 0.503 |
| path operator advantage | +0.075 [+0.048, +0.103] | +0.033 [+0.002, +0.062] |
| ratio of performance above chance | 2.21 | 2.04 |
| difference in advantage between strains: +0.042 [+0.002, +0.083] | | |
| wild type, 100 subsets of 18 recordings: +0.079 [+0.068, +0.088]; *unc-31* below all | | |
| wild type, largest 69% of responses: +0.075 | | |

Table S8: **Responses per stimulation, and detection at matched stimulation counts, in wild type and *unc-31* mutants.** Response occurrence is the fraction of stimulation–responder trials exceeding the 95th percentile of the responder's reference distribution (5% expected without a response). Interaction terms are wild type minus *unc-31*, per standard deviation of the predictor, with distance measured between somata; intervals are 95% bootstrap intervals over recordings. Matched detection uses the 12,520 pairs stimulated in both strains, with equal numbers of stimulations per pair, at the threshold |statistic| > 3.5.

| | wild type | *unc-31* |
|---|---|---|
| recordings with usable trials | 110 | 15 |
| stimulation–responder trials | 213,353 | 36,201 |
| released response rate, ≤ 5 stimulations | 2.0% | 2.4% |
| released response rate, > 5 stimulations | 6.2% | 4.0% |
| response occurrence, 1 step | 0.098 | 0.087 |
| response occurrence, 2 steps | 0.064 | 0.058 |
| response occurrence, 3 steps | 0.053 | 0.051 |
| response occurrence, ≥ 4 steps | 0.039 | 0.042 |
| matched detection rate | 4.7% | 4.7% |
| matched path operator AUC | 0.574 | 0.581 |
| matched path operator advantage | +0.032 [+0.004, +0.058] | +0.043 [+0.014, +0.078] |
| response occurrence, genotype × path operator: +0.002 [−0.003, +0.007] | | |
| response occurrence, genotype × distance: −0.001 [−0.008, +0.004] | | |
| response occurrence, genotype × direct synapse: +0.002 [−0.013, +0.016] | | |
| matched difference in advantage between strains: −0.011 [−0.046, +0.025] | | |

#### Degree-preserving surrogates with symmetric gap junctions

The main-text surrogates rewire the combined matrix as a directed graph, so the two orientations of a gap junction move independently. In a second set of 100 surrogates, chemical synapses were rewired by directed swaps and gap junctions by undirected swaps ($\{a, b\}, \{c, d\} \rightarrow \{a, d\}, \{c, b\}$, ten attempts per junction), and the two were summed. Each neuron kept its chemical in- and out-degree and its number of gap-junction partners, and gap junctions stayed symmetric.

Both sets of surrogates gave the same result (Table S9).

Table S9: **Degree-preserving surrogates.** AUC of the path operator for signal propagation on the real connectome and on 100 surrogates of each kind (mean ± s.d.); $z$ is the real value minus the surrogate mean, divided by the surrogate s.d.

| surrogates | pairs | real | surrogates | $z$ |
|---|---|---|---|---|
| combined matrix rewired as a directed graph | all observed | 0.637 | $0.522 \pm 0.007$ | 16.5 |
| | unsynapsed | 0.603 | $0.517 \pm 0.007$ | 11.4 |
| contact types rewired separately, | all observed | 0.637 | $0.523 \pm 0.008$ | 15.0 |
| gap junctions kept symmetric | unsynapsed | 0.603 | $0.517 \pm 0.008$ | 10.2 |

#### Chemical synapses and gap junctions separately

The path and class-level operators were rebuilt from chemical synapses alone (3,638 connections, 20,589 contacts) and from gap junctions alone (2,175 connections, 8,669 contacts, symmetrized by taking the larger orientation), each on all 302 nodes and normalized by its own spectral radius. Unsynapsed pairs were defined from the combined matrix throughout.

For each contact type the path operator outperformed direct connection strength by a similar margin, and cell classes kept all of its performance while random partitions kept less than a third (Table S10).

Table S10: **Chemical synapses and gap junctions separately.** Performance above chance (AUC − 0.5, or Spearman $\rho$) and fraction kept by cell classes, with 95% cluster-bootstrap intervals (500 resamples). Random: mean of 200 size-matched partitions.

| | contacts | direct | path operator | path − direct | cell classes kept (random) |
|---|---|---|---|---|---|
| signal propagation, | both | 0.062 | 0.137 [0.104, 0.171] | 0.075 [0.049, 0.103] | 1.00 [0.94, 1.08] (0.30) |
| all pairs | chemical | 0.036 | 0.108 [0.074, 0.142] | 0.072 [0.043, 0.102] | 1.16 [1.03, 1.33] (0.31) |
| | gap junction | 0.037 | 0.134 [0.104, 0.168] | 0.097 [0.072, 0.124] | 1.13 [1.03, 1.25] (0.23) |
| signal propagation, | both | – | 0.103 [0.063, 0.138] | – | 1.03 [0.91, 1.18] (0.07) |
| unsynapsed | chemical | – | 0.079 [0.042, 0.116] | – | 1.21 [1.05, 1.52] (0.14) |
| | gap junction | – | 0.104 [0.070, 0.135] | – | 1.17 [1.03, 1.42] (0.03) |
| activity correlation | both | 0.135 | 0.273 [0.235, 0.315] | 0.139 [0.111, 0.171] | 1.09 [1.02, 1.17] (0.26) |
| | chemical | 0.117 | 0.255 [0.215, 0.297] | 0.138 [0.104, 0.172] | 1.11 [1.03, 1.23] (0.27) |
| | gap junction | 0.098 | 0.248 [0.208, 0.289] | 0.150 [0.115, 0.190] | 1.09 [0.98, 1.23] (0.16) |

#### Predictive transfer at other merge counts

The main text reports 70 pairs among 161 candidates. The analysis was also run with 40, 55 and 75 pairs, and with the degree floor (86 candidates, 30 pairs); neither choice was fixed in advance.

Without the floor, L1-inferred partitions lost more than adult-inferred and less than random partitions at every number of pairs, but the separation from random was 1–1.6 s.d. at 40 and 55 pairs against 2.8–2.9 at 70 and 75. Thinning the adult to developmental synapse counts matched the L1 loss at 70 and 75 pairs but not at 40 or 55 (Table S11).

Table S11: **Predictive transfer at other merge counts.** Loss of adult signal-propagation prediction (% of the full operator's performance above chance). L1: mean (range) of the four L1 animals. Thinned adult: five replicates per animal, averaged. Cell classes and random: means of 50 partitions each. Last column: L1 minus random mean, in random-partition s.d. Bold: main text.

| candidates | pairs | adult | L1 (range) | thinned adult | cell classes | random | L1 vs random (SD) |
|---|---|---|---|---|---|---|---|
| 86 (floor) | 30 | 4.1 | 4.5 (3.1–5.4) | 4.0 | 0.7 | 7.6 | 0.9 |
| 161 | 40 | 0.2 | 9.5 (7.3–12.1) | 1.9 | −0.5 | 14.1 | 1.0 |
| 161 | 55 | −2.9 | 10.3 (6.6–12.7) | 5.4 | −0.8 | 18.1 | 1.6 |
| **161** | **70** | **2.4** | **7.4 (3.8–10.6)** | **7.1** | **−1.0** | **21.6** | **2.9** |
| 161 | 75 | 1.5 | 8.4 (5.4–12.0) | 8.1 | −1.3 | 22.0 | 2.8 |

#### Developmental analyses without glia

Repeating the developmental analyses without the four CEP sheath glia and the GLR cell, none of which met the degree floor used for merge selection, left every conclusion unchanged: transfer retained 92.3% rather than 90.8% of the within-animal advantage, and the thinned structural advantage of cell classes still showed no trend with developmental stage.

#### Lineage-only cell classes

Some cell classes were defined in part from connectivity, so the sufficiency of cell classes could partly reflect that definition. Lineage-only cell classes remove this possibility: left–right homolog pairs and the eight serially repeated ventral-cord families, with every other neuron in its own class ($k = 137$, a 2.2-fold reduction). Every lineage-only class lies within a single cell class. The class-level path operator on this partition kept all of the full operator's performance for every observable (Table S12), including signal propagation among unsynapsed pairs (1.14 [1.03, 1.35] of full performance), whereas size-matched random partitions kept 0.08. Left–right pairs alone ($k = 204$) gave the same result. The partition built without any connectivity-informed class definition performed at least as well as the cell classes, so connectivity entering historical class definitions does not account for their sufficiency.

Table S12: **Lineage-only cell classes.** Performance of the class-level path operator as a fraction of the full path operator's performance above chance, with 95% cluster-bootstrap intervals (500 resamples), against the mean of 200 size-matched random partitions. Response magnitude is scored over only 1,131 detected responses.

| | cell classes $k = 119$ | lineage-only $k = 137$ | left–right pairs $k = 204$ |
|---|---|---|---|
| signal propagation, all pairs | 1.00 [0.94, 1.08] | 1.07 [1.00, 1.15] | 1.07 [1.01, 1.16] |
| random | 0.30 | 0.34 | 0.60 |
| signal propagation, unsynapsed | 1.03 [0.91, 1.18] | 1.14 [1.03, 1.35] | 1.14 [1.03, 1.33] |
| random | 0.07 | 0.08 | 0.35 |
| activity correlation | 1.09 [1.02, 1.17] | 1.05 [1.01, 1.10] | 1.09 [1.04, 1.14] |
| random | 0.26 | 0.31 | 0.56 |
| response amplitude | 1.00 [0.90, 1.11] | 1.13 [1.06, 1.25] | 1.11 [1.04, 1.21] |
| random | 0.19 | 0.23 | 0.48 |
| response magnitude | 0.94 [0.71, 1.33] | 1.05 [0.87, 1.38] | 1.06 [0.87, 1.42] |
| random | 0.33 | 0.41 | 0.65 |

#### Activity correlations after removing the evoked response

Activity correlations and detected responses are computed from the same imaging frames, so shared evoked activity could make cell classes appear sufficient for both. Subtracting each neuron's stimulus-triggered template at every stimulation onset removes the stimulus-locked component while keeping every frame. On the residual traces, the full path operator predicted activity correlations at $\rho = 0.270$, the class-level path operator at $\rho = 0.295$, and size-matched random partitions at $\rho = 0.070 \pm 0.025$ ($z = 8.9$; 13,849 pairs). These values match those from the unaltered traces ($\rho = 0.273$, 0.297 and $0.072 \pm 0.025$; $z = 8.9$). Cell-class sufficiency for activity correlations therefore does not depend on the evoked component that the two observables share.

#### Coarse-graining by other grouping methods

Operator-defined classes were compared with three other ways of grouping neurons at the same resolutions (Table S13). The three operator-defined methods showed no detectable loss of signal-propagation or activity-correlation prediction down to $k = 9$, and sustained loss began at $k = 8$ for loss-guided agglomeration and $k$-means and at $k = 7$ for Ward clustering. $k$-means on direct connectivity profiles lost 35–47% of signal-propagation performance and 35–60% of activity-correlation performance at every resolution from $k = 20$ down. Spectral clustering lost 19% and 32% of signal-propagation performance at $k = 20$ and 15. Spatial clusters lost 25–38% of activity-correlation performance from $k = 20$ and 60–65% of response-amplitude performance at $k = 9$–20, while their loss of signal-propagation performance was not detectable until $k = 8$. The largest block of tied class-level operator values where loss first became detectable was 8–16% for the operator-defined methods, so the boundary is not produced by ties. What sets how far coarse-graining can go is therefore how neurons are grouped, and grouping by path-operator profile goes furthest.

Table S13: **Predictive loss by grouping method.** Loss as a fraction of the full path operator's performance above chance (0 = no loss, 1 = all lost). For $k$-means and spatial and spectral clustering, the mean over five initializations. * Detectable: lower bound of the 95% cluster-bootstrap interval (300 resamples, first initialization) above zero. Random: mean of 20 partitions size-matched to loss-guided agglomeration.

| $k$ | loss-guided (operator) | Ward (operator) | $k$-means (operator) | $k$-means (connectivity) | spectral | spatial | random |
|---|---|---|---|---|---|---|---|
| *signal propagation, all observed pairs* | | | | | | | |
| 20 | 0.02 | −0.11 | 0.01 | 0.36* | 0.19* | 0.10 | 0.96 |
| 15 | 0.00 | −0.04 | −0.07 | 0.35* | 0.32* | 0.11 | 1.00 |
| 12 | 0.08 | −0.04 | 0.00 | 0.43* | 0.11 | 0.19 | 1.03 |
| 10 | 0.09 | −0.09 | 0.05 | 0.38* | 0.13 | 0.22 | 0.98 |
| 9 | 0.09 | −0.08 | 0.07 | 0.47* | 0.12 | 0.22 | 0.99 |
| 8 | 0.23* | 0.13 | 0.03 | 0.44* | 0.13 | 0.20* | 1.00 |
| 7 | 0.28* | 0.19* | 0.06 | 0.51* | 0.35* | 0.28 | 0.98 |
| 6 | 0.19* | 0.38* | 0.16 | 0.52* | 0.30* | 0.56* | 1.08 |
| *activity correlations* | | | | | | | |
| 20 | −0.01 | −0.08 | 0.01 | 0.35* | 0.09 | 0.25* | 0.94 |
| 15 | −0.01 | −0.06 | −0.09 | 0.35* | 0.17* | 0.24* | 0.98 |
| 12 | 0.04 | −0.06 | 0.03 | 0.63* | 0.07 | 0.31* | 1.03 |
| 10 | 0.04 | −0.08 | 0.14 | 0.44* | 0.05 | 0.38* | 0.98 |
| 9 | 0.04 | −0.07 | 0.16 | 0.60* | 0.08 | 0.38* | 1.00 |
| 8 | 0.10* | 0.06 | 0.14* | 0.56* | 0.10 | 0.40* | 0.97 |
| 7 | 0.09 | 0.07 | 0.15* | 0.69* | 0.06 | 0.48* | 0.98 |
| 6 | 0.09 | 0.30* | 0.32* | 0.78* | 0.07 | 0.80* | 1.02 |

#### Comparison with anatomical and signaling communities

The transcribed memberships reproduce the agreement Dvali et al. report between their anatomical and signaling communities (adjusted Rand index 0.105; 0.060 without the pharynx). On the same 188 head neurons, the nine operator-defined classes agreed with the anatomical communities at 0.31–0.36 and with the signaling communities at 0.12–0.14 (Table S14). Without the pharynx these fell to 0.25–0.28 and 0.07–0.09. Operator-defined classes refit on the head neurons alone at $k = 20$, the resolution of the anatomical communities, agreed with them at 0.46–0.58 and with the signaling communities at 0.11–0.12. Every agreement exceeded all 2,000 permutations of class membership. Agreement with the signaling communities was therefore no greater than the agreement between Dvali et al.'s two partitions. The a priori subsystems, which use no connectivity, agreed with the signaling communities more than any connectivity-derived partition did (0.18). Where the operator-defined classes and the anatomical communities agree, the correspondence is often complete. All 20 neurons of the pharyngeal anatomical community fall in one loss-guided class, which also contains 15 of the 17 neurons of the pharyngeal signaling community, and the anatomical communities of labial and cephalic neurons (IL1D/V, IL2V, RIP, URA) and of head-proximal ventral-cord motor neurons map 12 of 12 and 11 of 13 neurons into single classes.

Coarse-graining the head into the anatomical communities, with every other neuron kept in its own class, produced no detectable loss of the path operator's prediction of any observable (Table S15), whereas random partitions with the same class sizes kept at most 0.09 of full performance. The operator-defined classes, restricted to the head in the same way, were also sufficient. With pairs involving pharyngeal neurons excluded, the anatomical communities kept 1.04 of signal-propagation performance and 1.23 among unsynapsed pairs, against 0.69–0.92 and 0.81–0.92 for the nine operator-defined classes and 0.72–0.84 and 0.59–0.81 for operator-defined classes refit on the head at $k = 20$. Within the head outside the pharynx, the anatomical communities therefore retain at least as much of the connectome's prediction as operator-defined classes of the same or coarser resolution.

Table S14: **Agreement with Dvali et al. communities.** Adjusted Rand index on the 188 head neurons and on the 168 remaining after the pharyngeal anatomical community is excluded. Operator-defined classes were built on the full connectome and restricted to the head, except those marked head-fit. $k$-means values are means over five initializations.

| | anatomical (20) | | signaling (6) | |
|---|---|---|---|---|
| | head | no pharynx | head | no pharynx |
| loss-guided agglomeration, $k = 9$ | 0.36 | 0.27 | 0.12 | 0.07 |
| Ward on operator, $k = 9$ | 0.35 | 0.28 | 0.14 | 0.09 |
| $k$-means on operator, $k = 9$ | 0.31 | 0.25 | 0.12 | 0.08 |
| Ward on operator, head-fit $k = 20$ | 0.58 | 0.48 | 0.12 | 0.07 |
| $k$-means on operator, head-fit $k = 20$ | 0.46 | 0.36 | 0.11 | 0.06 |
| subsystems (a priori) | 0.35 | 0.27 | 0.18 | 0.14 |
| cell classes | 0.21 | 0.23 | 0.03 | 0.03 |
| Dvali et al. signaling communities | 0.105 | 0.060 | – | – |

Table S15: **Class-level path operator on the head.** Head neurons grouped by each partition, every other neuron in its own class. Performance on pairs with both neurons in the head as a fraction of the full path operator's on the same pairs, with 95% cluster-bootstrap intervals (300 resamples) for single partitions and the spread across initializations for $k$-means. Random: 50 partitions with the anatomical community sizes.

| | signal propagation (all pairs) | unsynapsed pairs | activity correlation | response amplitude |
|---|---|---|---|---|
| anatomical communities (20) | 1.17 [0.98, 1.48] | 1.31 [1.05, 1.92] | 1.09 [0.99, 1.19] | 1.18 [1.00, 1.45] |
| loss-guided, $k = 9$ | 0.94 [0.78, 1.16] | 1.09 [0.85, 1.49] | 0.97 [0.87, 1.06] | 0.98 [0.86, 1.14] |
| Ward on operator, $k = 9$ | 1.06 [0.90, 1.32] | 1.12 [0.83, 1.65] | 1.09 [0.99, 1.19] | 1.17 [0.97, 1.42] |
| $k$-means on operator, $k = 9$ | 0.91 [0.69, 1.03] | 0.93 [0.64, 1.10] | 0.92 [0.78, 0.98] | 0.99 [0.77, 1.14] |
| Ward on operator, head-fit $k = 20$ | 1.07 [0.90, 1.36] | 1.14 [0.87, 1.66] | 1.07 [0.98, 1.15] | 1.13 [0.97, 1.36] |
| cell classes | 0.99 [0.92, 1.07] | 1.00 [0.90, 1.12] | 1.12 [1.05, 1.20] | 0.98 [0.89, 1.09] |
| random | 0.08 | 0.04 | 0.09 | −0.03 |

#### Stimulus excision, sham excision and detrending

Removing the volumes from two before to $w$ after each stimulation left cell-class sufficiency for activity correlations unchanged at every window up to $w = 40$ volumes (Table S16). The full and class-level path operators both declined only at $w = 60$, where almost the entire interval between stimuli (67.9 volumes on average) is removed, and the class-level operator still exceeded the full operator. A sham that removed the same number of windows of 42 volumes at random onsets reproduced true excision closely across three seeds (class-level $\rho = 0.285$–$0.296$ against 0.291 for true excision; $z = 8.7$–$8.9$ against 8.4). Per-neuron linear detrending, alone or combined with 20-volume excision, also left the result unchanged. As described in the Supplementary Methods, the evoked transient outlasts any excision window, so excision is reported as a check that correlation sufficiency does not depend on particular frames; the stimulus-template subtraction above is the test that addresses shared evoked activity directly.

Table S16: **Activity correlations under excision, sham excision and detrending.** Spearman $\rho$ between the symmetrized predictor and activity correlation over 13,849 pairs. Random: mean of size-matched random partitions; $z$ compares the class-level operator with them.

| treatment | full operator | class-level (cell classes) | random | $z$ |
|---|---|---|---|---|
| none | 0.273 | 0.297 | 0.072 | 8.9 |
| excise 10 volumes | 0.275 | 0.299 | 0.072 | 8.9 |
| excise 20 volumes | 0.275 | 0.299 | 0.072 | 8.8 |
| excise 30 volumes | 0.269 | 0.294 | 0.070 | 8.5 |
| excise 40 volumes | 0.265 | 0.291 | 0.069 | 8.4 |
| excise 60 volumes | 0.180 | 0.199 | 0.043 | 6.3 |
| sham, 40 volumes (3 seeds) | 0.262–0.273 | 0.285–0.296 | 0.069–0.070 | 8.7–8.9 |
| detrended | 0.266 | 0.301 | 0.067 | 9.5 |
| detrended, excise 20 volumes | 0.265 | 0.300 | 0.067 | 9.4 |
| stimulus template subtracted | 0.270 | 0.295 | 0.070 | 8.9 |

#### Calibration of $\gamma$

At $n = 302$ and the cell-class size distribution, a planted, exactly equitable structure gives $\gamma = 0$, the same structure with 50% within-class noise gives $0.233 \pm 0.003$, and matrices without class structure give $0.778 \pm 0.002$ (Gaussian) and $0.760 \pm 0.008$ (shuffled connectome). The cell classes on the connectome give $\gamma = 0.300$ (outgoing 0.287, incoming 0.312), against $0.755 \pm 0.014$ for size-matched random partitions of the same connectome ($z = -33$). The cell classes are therefore not exactly equitable, but they depart from equitability about as much as a planted structure with within-class noise at half the block value, and far less than any partition without class structure.

#### Degree floors for the merge criterion

Without a degree floor, the merge criterion appeared stronger than it is (Table S17). Forty of its 60 selected pairs joined two neurons below median degree, and such neurons have nearly empty, and therefore similar, connectivity profiles. At the median floor, no selected pair contained a below-median neuron, and the criterion still lost less predictive performance among unsynapsed pairs than random disjoint pairs (+0.0054 against +0.0164) and much less structure (0.065 against 0.175), with 38% of merges joining two neurons of the same cell class against a chance rate of 0.9%. About a third of the unfiltered advantage over random pairs therefore came from low-degree neurons. At a floor of 40, only 83 neurons and 41 disjoint merges remain, and random merges among well-connected neurons also become harmless (+0.0171), so that row loses the contrast needed to test the criterion rather than showing that it fails.

Table S17: **Merge criterion under degree floors.** Merges among neurons at or above each total-degree floor. Predictive loss: AUC lost among unsynapsed pairs (full path operator AUC $-$ 0.5 = 0.1026). Structural loss over all ordered pairs. Random: mean $\pm$ s.d. of 30 sets of random disjoint pairs from the same neurons. Same-class rate with its chance rate in brackets, the probability that two distinct neurons available at that floor belong to the same cell class.

| degree floor | none | 20 | 29 (median) | 40 |
|---|---|---|---|---|
| cells available | 302 | 235 | 152 | 83 |
| merges | 60 | 60 | 60 | 41 |
| predictive loss, merge criterion | −0.0078 | +0.0062 | +0.0054 | +0.0162 |
| predictive loss, left–right pairs | −0.0136 | −0.0046 | −0.0057 | +0.0009 |
| predictive loss, random | $+0.045 \pm 0.016$ | $+0.042 \pm 0.011$ | $+0.016 \pm 0.010$ | $+0.017 \pm 0.004$ |
| structural loss, merge criterion | 0.056 | 0.083 | 0.065 | 0.046 |
| structural loss, left–right pairs | 0.037 | 0.034 | 0.026 | 0.016 |
| structural loss, random | $0.303 \pm 0.035$ | $0.248 \pm 0.022$ | $0.175 \pm 0.011$ | $0.129 \pm 0.006$ |
| same-class rate (chance) | 47% (1.1%) | 30% (1.0%) | 38% (0.9%) | 44% (1.2%) |
| pairs with both neurons below median degree | 40/60 | 18/60 | 0/60 | 0/41 |
